# Slitrk/LAR-RPTP and disease-associated variants control neuronal migration in the developing mouse cortex independently of synaptic organizer activity

**DOI:** 10.1101/2023.10.20.563293

**Authors:** Vera P. Medvedeva, Pierre Billuart, Alice Jeanmart, Lisa Vigier, Jaewon Ko, Lydia Danglot, Alessandra Pierani

**Author notes:** Corresponding author: Alessandra Pierani, Institute of PSYCHIATRY AND NEUROSCIENCE of Paris, INSERM UMR 1266, 102-108 Rue de la Santé, 75014 PARIS, FRANCE.

## Abstract

Slitrks and their ligands LAR-RPTPs are type I transmembrane proteins previously implicated in the etiology of various neuropsychiatric disorders including obsessive-compulsive disorders (OCDs) and schizophrenia. Over the last decade, their functions were extensively studied in hippocampal neurons *in vitro* and shown to shape synapse organization. Although both protein families are highly expressed prior to synapse formation, their function in earlier steps of cerebral cortex development remains unknown. Here we investigated the role of Slitrk1, Slitrk2, Slitrk3 and LAR-RPTPs (Ptprs and Ptprd) in the embryonic mouse cortex by acute genetic manipulation using *in utero* electroporation. All genes, except Slitrk3, promoted specific alterations in radial migration of glutamatergic neurons. Slitrk1 and Slitrk2 overexpression was associated with accumulation of neurons in distinct regions of the cortical plate. Using deletion mutants and a series of Slitrk variants associated with neurodevelopmental disorders (NDDs), we showed that distinct domains are crucial for intracellular Slitrk1 distribution and/or density and shape of VAMP2^+^ presynaptic boutons. Interestingly, bouton alterations did not correlate with the observed migration delays, suggesting that Slitrk1 influence cell migration independently on its synaptogenic function. Furthermore, co-electroporation experiments with LAR-RPTPs, mimicking their co-expression observed by scRNAseq, rescued the migration deficits, suggesting possible *cis*-interactions between Slitrks and LAR-RPTPs. Together, these data indicate that in the embryonic cerebral cortex Slitrks and LAR-RPTPs cooperate in consecutive steps of radial migration through distinct mechanisms than in synapse organization and support a relevant role of Slitrk/LAR-RPTP dysfunctions in NDDs at earlier stages of cortical development.

## Introduction

Slitrks (SLIT and NTRK-like) and LAR-RPTPs (Leukocyte Common Antigen-related Receptor Protein Tyrosine Phosphatases) are transmembrane protein families, which have recently emerged as key synaptic organizers [1–3]. The Slitrk family comprises 6 paralogs (Slitrk1-6) with two extracellular domains enriched with leucine rich repeats (LRR), namely LRR1 and LRR2, and intracellular regions bearing various number of potential tyrosine phosphorylation sites [4]. PTPR type S or type D (Ptprd and Ptprs, also known as Ptprσ and – δ), belongs to the type IIa Receptor-like Protein Tyrosine Phosphatase (RPTP) family [5]. They interact with Slitrks through extracellular immunoglobulin-like domains [3]. It was previously reported that Slitrk LRR1 mediates the interactions with LAR-RPTPs [6–8]. LAR-RPTPs were shown to require the inclusion of the MeB microexon within their ECD for interaction with Slitrks and to organize the synaptic assembly [6]. *In vitro* studies have shown that postsynaptic Slitrks are involved in the formation of both excitatory and inhibitory synapses through interactions with presynaptic Ptprs and Ptprd, respectively, with an exception for Slitrk3, which is exclusively involved in inhibitory synaptogenesis through interaction with Ptptrd [9]. The synaptic functions of Slitrk2, Slitrk3 and Slitrk5 have been described in distinct brain regions [10–13]. However, it remains unclear whether known synaptic functions of Slitrks rely on their interactions with LAR-RPTPs *in vivo*.

Both SLITRKs and LAR-RPTPs have been associated with a range of neuropsychiatric disorders, including obsessive compulsive spectrum disorders (OCDs) (for *SLITRK1* and *PTPRD*), Gilles de la Tourette’s syndrome (for *SLITRK1, SLITRK3, SLITRK5* and *SLITRK6*), autism spectrum disorders (ASDs) (for *SLITRK3*, *PTPRD* and *PTPRS*), schizophrenia (SCZ) (for *SLITRK1*, *SLITRK2* and *SLITRK4*), addictive behaviors and cognitive impairments (for *PTPRD*) [2, 3, 14–18]. They are vastly expressed in the nervous system and besides synapse organization Slirtk1 and Slirtk2 are also reported to regulate neuritogenesis *in vitro* [4, 19, 20]. In addition, expression of both protein families is detected during early corticogenesis in mice, before synaptic formation, and each gene exhibits a distinct expression pattern [4, 21–23].

To investigate whether Slitrks and LAR-RPTPs perform additional functions during glutamatergic neuron differentiation prior to functional synapse formation, we overexpressed (OE) Slitrks or disease-associated mutants in the embryonic cortex using *in utero* electroporation (IUE). We revealed that they alter specific steps of radial migration, synapse formation and intracellular protein distribution through distinct protein domains. Together, our data show that Slitrks and LAR-RPTPs perform specific functions through both *trans*-and possibly *cis*-interactions in multipolar migration and glia-mediated locomotion as well as in the control of synaptic density and vesicle docking at presynaptic sites.

## Materials, Patients and Methods

### Experimental Design and Statistical Analysis

For all experiments cortex from mouse embryos were electroporated using plasmid vectors to express Slitrk or Ptprs proteins or variants and compared their distribution to either empty vector expressing GFP alone or to wild-type proteins when addressing the variants consequences.

Quantitative data are expressed as the group mean and standard deviation, unless stated otherwise in the figure legend. For quantifications at least 3 embryos from at least 2 different litters per group were used, and up to 3 sections per embryo for analysis were selected. The total number of sections in the groups are indicated as n= in the figure legends.

For the statistical analyzes datasets normality were tested using by the Shapiro-Wilk normality test. For direct comparisons, *F* test was used for homogeneity of variance and the data were analyzed using t-test or Mann-Whitney test (U value is reported). For multiple comparisons, one-way ANOVA (F values) or Kruskal-Wallis test (KW value and Dunn’s multiple comparisons test are reported) for non-parametric comparisons, or two-way ANOVA (F values for Interaction statistics is reported and Sidak’s multiple comparisons test, unless otherwise specified) or Brown-Forsythe ANOVA, unpaired t with Welch’s correction for multiple comparisons were performed and mentioned in figure legends. All statistical analyses and graphical representations were performed using Prism (GraphPad Prism 9.00), and statistical significance was indicated by *P< 0.05, **P< 0.001, ***P< 0.001, ****P< 0.0001 or not significant (n.s.) for P ≥ 0.05.

### Animals

All animals were handled in accordance with good animal practice as defined by the national animal welfare bodies, and all mouse work was approved by the Ministry of Higher Education, Research and Innovation, France (Ministere de l’Enseignement supérieur, de la Recherche et de l’Innovation) and the Animal Experimentation Ethical Committee of Paris Descartes University (CEEA-34) (Reference: 2018012612027541).

Swiss time-pregnant mice were purchased at Janvier Labs, France and housed under a 12 h light/dark cycle with standard diet and water *ad libitum*. For timed mating, noon of the day of vaginal plug detection was designated as E0.5. Both male and female embryos were used for the experiments, without distinction.

### In Utero Electroporation and DNA plasmids

Mouse embryos were electroporated at E13.5 as previously described [24, 25]. The five mouse genes ORFs, were cloned into the polylinker (LinkerA) site of a pCAGGS-LinkerA-*ires*-nls-GFP backbone vector (pCAG). The following isoforms were used: Slitrk1 ^(^NM_199065.2), Slitrk2 (NM_198863.2), Slitrk3 (NM_198864.3), Ptprs (NM_001252453.1), Ptprd (NM_001352630.1). Slitrk1 intra-cellular (ICD amino acid positions 647-696) or extra-cellular (ECD amino acid positions 23-618) domains were substituted with a triple HA sequence. Slitrk1 and Slitrk2 human variants were introduced into the mouse sequence by oligonucleotide directed mutagenesis at homologous positions using an In-Fusion HD Cloning Plus kit (Takara Bio Europe).

### Tissue preparation, in situ hybridization and immunohistochemistry

Embryo brains were recovered at E17.5 and processed as previously described [24, 25]. Immunohistochemistry and *In situ* hybridization experiments were performed on 16 µm or 20 µm thick Coronal sections, respectively [26]. We used the following primary antibodies: goat anti-Slitrk1 (AF3009, R&D Systems, 1:200), rat anti-Ctip2 (ab18465, Abcam, 1:300), goat anti-Brn2 (sc-6029, Santa Cruz Biotechnology, 1:250), rabbit anti-pH3 (06-570, Merck, 1:500), rabbit anti-Pax6 (PRB-278P, BioLegend, 1:500), rabbit anti-Tbr2 (ab216870, Abcam, 1:2000), goat anti-CTGF (sc-14939, Santa Cruz Biotechnology, 1:250), mouse anti-Tuj1 (MAB1195, R&D Systems, 1:20), rabbit anti-Tbr1 (ab31940, Abcam, 1:500), rabbit anti-Foxp2 (ab16046, Abcam, 1:500), rabbit anti-VAMP2 (104202, Synaptic Systems, 1:500), mouse anti-HA.11 (16B12, Covance, 1:1000), chicken anti-GFP (GFP-1020, Aves Labs, 1:1000), rabbit anti-cleaved Caspase-3 (Cell Signaling, 1:400).

### Image acquisition and analysis

Images for evaluation of migration delay, cell fate markers and morphology screening were all acquired using a NanoZoomer scanner (Hamamatsu, Japan). After conversion into Tiff format with ImageJ plugin NDPI tools, images were analyzed using the ICY bioimaging software [27].

For quantification of migration delay in the CP we divided the electroporated site into 5 equal bins and created 5 rectangular ROI to quantify the number or the intensity of GFP+ cells. The CP depth was defined by DAPI staining density, with an upper and lower borders defined as the edges of upper and lower nuclei dense layers, thus low density layers, the marginal zone (MZ) and the subplate (SP), were omitted in this analysis. Quantifications of cells in the intermediate zone (IZ) were always performed by cell somas counting under the CP (under the DAPI nuclei dense lower border). Quantifications were performed in the lateral cortex along 3 coronal coordinates (when electroporation was present) per embryo: GD18 cor. 9, cor. 12 and cor. 15 (Atlas of the prenatal mouse brain. Schambra, Lauder, Silver. 1992), or in the dorsal cortex (only Slitrk3 OE confirmation). Ramified ectopic cells were counted throughout all the rostro-caudal levels of lateral cortices where electroporations were present. For more detailed cellular level analysis, we used a Leica (SP8) confocal microscope and 20x, 40x and 100x magnifications. Slitrk1 OE neuronal identity quantifications, Slitrks soma length and dendrite travel distances, Slitrk1 and mutant OE proteins cellular distribution as well as VAMP2 analysis were performed using the ICY bioimaging software.

### VAMP2 positive presynaptic boutons analysis

Brain stacks images were acquired with a confocal laser scanning microscope (LEICA SP8 STED 3DX) equipped with a 93×/1.3 NA glycerol objective, a white-light laser and hybrid detectors so that images of 1600×1600 pixels were acquired with a pixel size in the range of 80 nm.

VAMP2 clusters in proximal contact with GFP cells were analyzed using a dedicated program coded within Icy software and “protocol”. VAMP2 clusters were segmented in 3D using Icy spot detector plugin with wavelet analysis as described previously [28]. Diameter of a synapse was evaluated previously by 3D scanning electron microscopy (FIB/SEM) on cortical layers [29] and is ranging from 350 to 450nm. Clusters with a surface area bigger than 3 pixels were considered in further analysis which likely recognizing presynaptic boutons of all the possible sizes (between 0.006μm^3^ and and 20μm^3^). GFP cells were automatically segmented using Hierarchical K-Means plugin after a 0,8 gaussian filter and a 4 classes segmentation. To estimate the region of synaptic junction touching GFP^+^ cells, we thus segmented the GFP^+^ cells in 3D (**Fig. S6C**) and slightly dilate the 3D envelope of a few hundreds nanometers to englobe touching presynaptic boutons (**Fig. S6E**). GFP cells envelope were then dilated of 3 pixels (240 nm), and intersecting VAMP2 clusters with this dilated contour were considered as proximal VAMP2 positive boutons. Morphological characteristics of 3D VAMP2 spots were analysed for size, intensity and spherical shape. Roundness is the normalised ratio between the radius of the minimum inscribed and largest circumscribed sphere expressed as a percentage (100% for a circle or sphere). The protocol indicates the roundness index (100% being a perfect sphere) and calculates mean staining intensity, thus providing an idea on enrichment of VAMP2 molecules in the bouton. All informations were exported automatically in excel files. We have observed that VAMP2 spots tend to align the GFP^+^ cell body margin inside and outside the detectable GFP signal, thus VAMP2 spots in GFP and contour (GFP*) are likely located to the presynaptic boutons contacting GFP^+^ cells. For the data collection in the CP we considered the MZ and underlying layers of 6-7 neuronal bodies (DAPI^+^) - roughly similar to CP1; for the IZ we took an area with at least 3 GFP^+^ cellular bodies in the field, when possible. Independent stacks (1600×1600pixels x 33 z slices) were acquired on independent brain slices within CP and IZ. All images were acquired with the same acquisition parameters and data were normalized to the control.

## Results

### Slitrk/LAR-RPTP functional domains and expression patterns in the mouse developing cerebral cortex

Possibly pathological SLITRK1 variants (black arrows) found in NDD patients are remarkably enriched outside LRR1, with a large prevalence to the LRR2 domain (**Fig. 1A,B**). Meanwhile, only two out of five *SLITRK2* variants (orange) are located within the LRR1 and LRR2 domains. This suggests new functions of domains outside LRR1 underlying SLITRK-related pathogenesis in NDDs.

To precisely map the spatial and temporal expression patterns of Slitrks/LAR-RPTPs in the developing cerebral cortex we mined biological databases at E14.5 (from the Genepaint *in situ* hybridization database, **Fig. 1C**) and from E13 to E17 (**Fig. S4A**, **S5** from the [Genebrowser](http://genebrowser.unige.ch/telagirdon/#query_the_atlas) scRNAseq [30]). We observed that Slitrk1 is expressed in the emerging CP (**Fig. 1C**) at E14.5 and throughout late embryogenesis in most neuronal populations (**Fig. S4A**). Unlike Slitrk1, Slitrk2 is mostly absent from early-born neuronal populations (**Fig. 1C** at E14.5 and **Fig. S4A** (E12-13-born analyzed at E16-17)) and is expressed in neuronal populations born at E14-15 and collected at E18.5-P0, distinct from those expressing Slitrk1. Indeed, the Slitrk1 and Slitrk2 expression patterns appeared to be complementary over time. Slitrk3 expression in the cortical plate (CP) is very low (**Fig. 1C** at E14.5 and **Fig. S5**) [4, 30]. Ptprs and Ptprd are strongly enriched in the postmitotic compartment (**Fig. 1C**) in all neurons born at different embryonic stages (**Fig. S4A**). However, they display stable over time but complementary expression patterns during early and late stages of neuronal differentiation (N1d and N4d, 24h and 96h after neuronal birthdate), with Ptprs expression higher in young and Ptprd in older neurons (**Fig. S4A**). The complementary expression of Slitrks and LAR-RPTPs suggests possible distinct and consecutive roles in glutamatergic neuron generation, migration and/or maturation.

**Figure 1.**
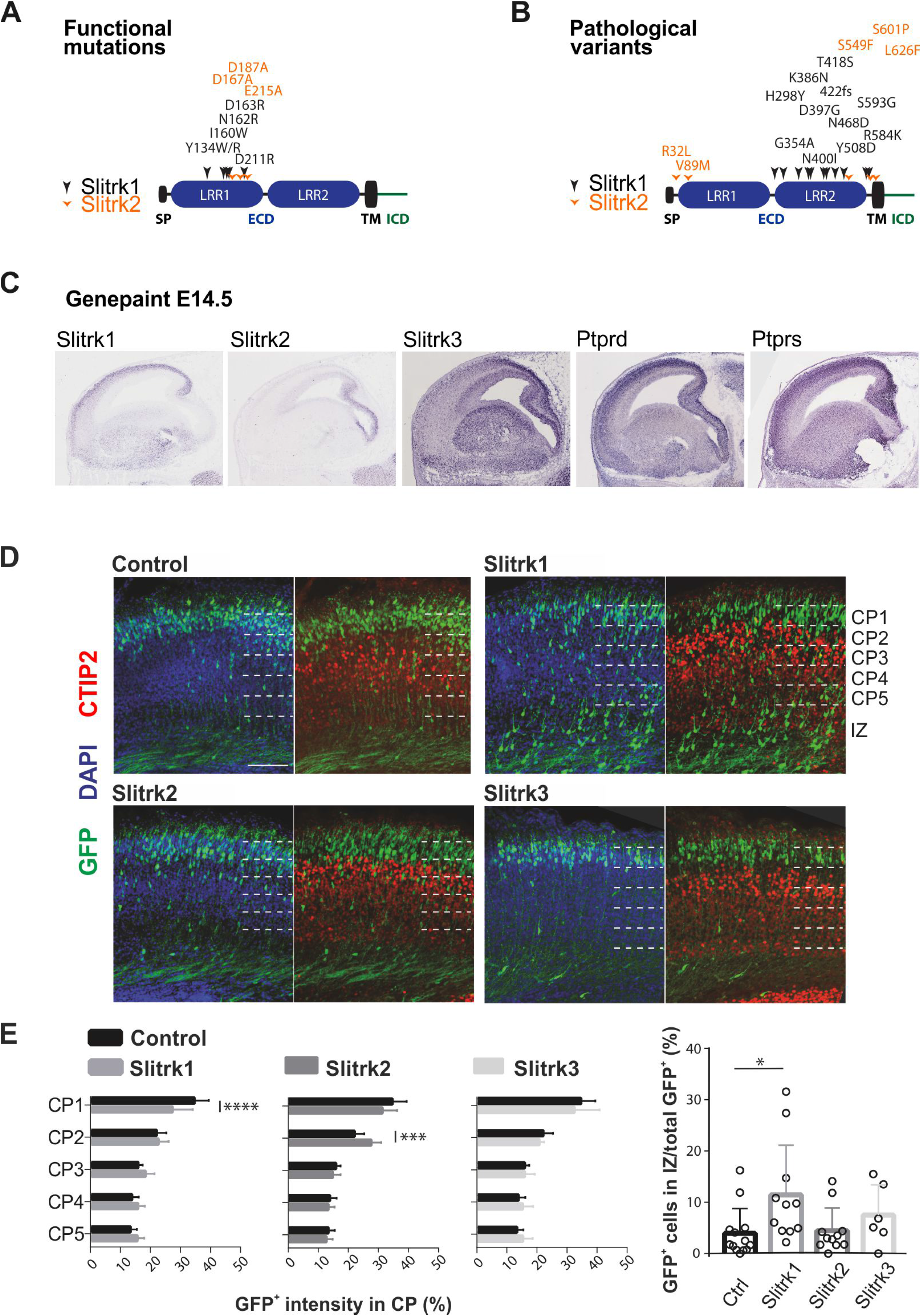
Structure-function analyses of disease-associated mutations in both *SLITRK1* and *SLITRK2*. Gene expression of Slitrks and Ptprs. Slitrk1 and Slitrk2 overexpression alters radial migration of glutamatergic neurons in the mouse embryonic cerebral cortex. **A)** Critical residues necessary for LAR-RPTPs-Slitrks interaction shown in *in vitro* studies [6–8]. LRR1 is the minimal domain required for synaptogenesis and binding with LAR-RPTPs proteins. **B**) Location of known *SLITRK1* and *SLITRK2* disease-associated variants leading to amino acid substitutions [14, 17, 31, 32, 34, 63, 64]. SP: signal peptide, TM: transmembrane domain, LRR: leucine rich repeat domain, ECD: extracellular domain, ICD: intracellular domain. **C**) LAR-RPTPs-Slitrks expression in the developing cortex. Sagittal view of E14.5 developing mouse cerebral cortex showing *in situ* hybridization data from the Genepaint database (https://gp3.mpg.de/). **D**) Representative immunofluorescence confocal images of cortical sections stained with DAPI (dense in the CP) and Ctip2 (marker of layer 5, 6 and the SP) highlighting distinct degrees of migration delays. Scale bar: 0.1mm. **E**) Quantification of migration delays throughout the CP divided in 5 equal bins and under the CP in the IZ. Only significant differences are marked in the CP (Control *vs* Slitrk1 *F*(4, 100)=8.174, *P*<0.0001; *vs* Slitrk2 *F*(4, 100)=6.342, *P*=0.0001; *vs* Slitrk3 *F*(4, 75)=0.9271, *P*=0.4529), and in the IZ (KW=9.775, *P*=0.0208). Each point represents 1 brain section (n=5-13).

### Overexpression of Slitrk1 and Slitrk2 affect neuronal migration during early neocortical development

To examine whether Slitrks could regulate cortical development *in vivo*, we overexpressed Slitrks proteins by *in utero* electroporation (IUE) in the lateral pallium (prospective neocortex) between E13.5 and E17.5. Glutamatergic neurons overexpressing Slitrk1 or Slitrk2 (**Fig. S1)** exhibited migration delays with distinct patterns, whereas Slitrk3 expression had no effect (**Fig. 1D, E**). Slitrk1 OE induced the most pronounced phenotype with a subpopulation of GFP^+^ neurons stalled underneath the CP within the intermediate zone (IZ) and the rest reaching the upper CP with a significant delay detectable in the most superficial part (bin 1 or CP1). The overall GFP comparable intensities in the CP, together with the absence of cell stagnation in the VZ or increase in activated-Caspase3 staining (**Fig. S2**), suggest that neither cell proliferation or death are affected by OE. Delayed cells in both the CP and the IZ exhibited a normal identity of glutamatergic neurons born at E13.5 (Brn2^+^/Ctip2^-^) (**Fig. S2**). Moreover, analysis of PH3, Pax6, Tbr2, Tuj1, Tbr1, CTGF and NURR1 indicated no gross alterations in cell-autonomous or non-autonomous identity in the electroporated areas compared to control side.

The Slitrk2 OE, although apparently less severe as it did not promote stalling in the IZ, produced a delay in the CP different from the Slitrk1 OE, with higher neuronal accumulation in the CP2 bin when compared to controls (**Fig. 1D,E**).

Overall, these data suggest that acute manipulation of Slitrk1 and Slitrk2 expression in the developing cortex induces specific defects in glutamatergic neurons migration and positioning.

### Slitrk1 and Slitrk2 domains and synaptogenic variants differentially influence migration

We next asked which Slitrk1 domains are involved in regulating migration. We replaced the intracellular (ICD) or extracellular (ECD) domain with triple HA-tag upon conservation of the signal peptide and transmembrane domain (TM) to ensure genuine subcellular targeting of the proteins (**Fig. 2A**). Deletion of either ICD (ΔICD) or ECD (ΔECD) prevented the migration delay observed with the full-length Slitrk1 (Slitrk1) in CP1. (**Fig. 2B, C**). In contrast, whereas deletion of the ECD has no effect in the IZ, Slitrk1 ΔICD still exhibited few GFP^+^ cells in the IZ (**Fig. 2B**). These results (**Table S1**) suggest that the function of Slitrk1 domains may not be equivalent in the CP and IZ.

**Figure 2.**
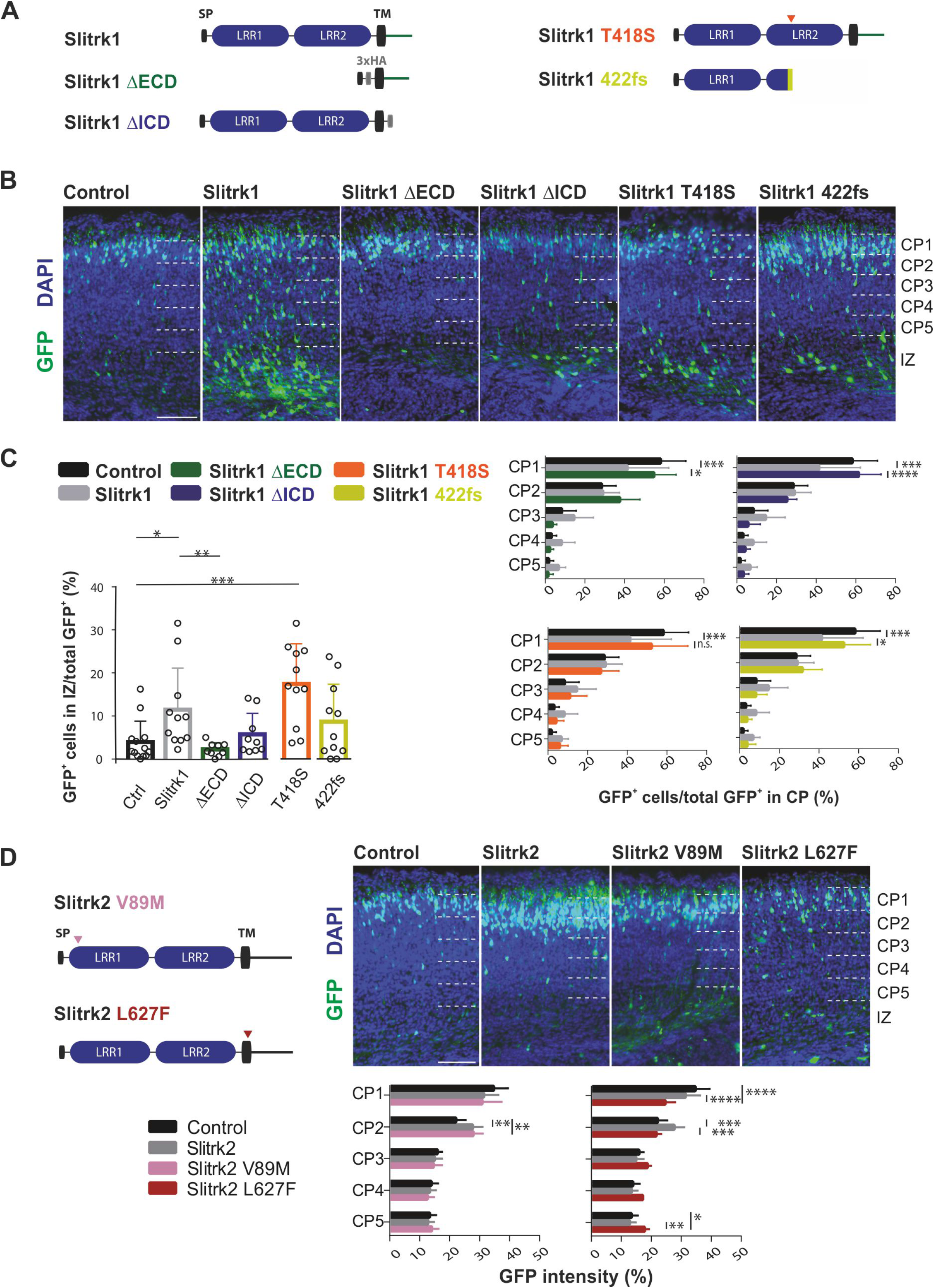
Effects of Slitrk1 or Slitrk2 mutations on neuronal migration in the CP and IZ. **A**) Schemas of Slitrk1 deletions and point mutations (orange arrowheads) and (**B**) representative immunofluorescence images at E17.5 in the lateral cortex upon their IUE at E13.5. **C**) Quantifications of migration delay in the CP, 5 equal bins (right graphs; Control *vs* Slitrk1 *vs* Slitrk1 ΔECD *F*(8, 145)=5.207, *P*<0.0001; *vs* Slitrk1-ΔICD *F*(8, 140) = 5.075, *P*<0.0001; *vs* Slitrk1 T418S *F*(8, 145) = 2.813, *P*=0.0063; *vs* Slitrk1 422fs *F*(8, 155) = 3.419, *P*=0.0012) and of cells present in the IZ (left graph; Control *vs* Slitrk1 vs Slitrk1 ΔECD and ΔICD KW=13.31, *P*=0.004; Slitrk1 *vs* Slitrk1-T418S *U*=41, *P*=0.2169; Slitrk1 *vs* Slitrk1-422fs *U*=47, *P*=0.3913). Only significant differences are marked. Each point represents 1 brain section (n=9-13). **D**). Slitrk2 mutations location schema and their representative immunofluorescence images at E17.5 upon IUE in the lateral cortex at E13.5 (top). Quantification of migration delays in 5 equal bins (below). Only significant differences are marked (Control *vs* Slitrk2 *vs* Slitrk2 V89M *F*(8, 125)=3.952, *P*=0.0003; *vs* Slitrk2 L627F *F*(8, 130)=14.35, *P*<0.0001), n=6-12. Scale bars: 0.1mm. 3xHA: triple HA tag (gray box), SP: signal peptide, TM: transmembrane domain, LRR: leucine rich repeat domain.

To investigate whether variants associated with “synaptopathies” could also affect Slitrk function in radial migration, we generated a series of Slitrk1 and Slitrk2 point mutants affecting either the LLR2 or the LRR1 domains, respectively. (**Fig. 2A and 2D**). Slitrk1-T418S is the most frequent point mutation associated with Tourette Syndrome and OCDs [17, 31, 32]. *In vitro* it affects both synaptogenesis and neurite outgrowth, and is expected to disrupt the 3D structure within the LRR2 domain [33]. However, in vivo, it behaved similarly to Slitrk1 in both the CP and the IZ (**Fig. 2B, C**), suggesting that this variant does not alter the Slitrk1 effects on migration delay and accumulation in the IZ. The second Slitrk1 mutation (422fs), associated with Tourette syndrome and Trichotillomania, leads to a large truncation of the protein leaving the LRR1 portion intact but lacking the TM and ICD domains [34] Similarly to the Slitrk1 ΔICD, the 422fs expression showed no effect on migration with no difference with respect to control in CP1 highlighting the importance of the ICD and/or the TM in mediating migration delays. Importantly, the “synaptogenic” LRR1 domain alone preserved in both T418S and 422fs variants is not sufficient to alter migration in both the CP and IZ.

We also examined two disease-associated Slitrk2 variants using IUE (**Fig. 2D**). Among Slitrk2 variants linked with SCZ, L626F (located in the TM domain with predicted damaging effects) presents with normal cell-surface expression in dendrites while the V89M mutation (located in the LRR1) impairs its dendritic targeting and the synapse formation in neurons while maintaining its synapse-inducing ability in co-culture assays [33]. The Slitrk2 V89M induced a migration delay similar to Slitrk2 WT (Slitrk2), while the Slitrk2 L627F (equivalent to the human L626F substitution) displayed a stronger effect with significantly less GFP^+^ cells in the upper CP (CP1 and CP2) and accumulation in the lower CP (CP5) (**Fig. 2D**). Thus, a variant previously known to strongly alter presynaptic organization [33] does not appear to influence the Slitrk2 effect on migration, while a predicted damaging mutation in the TM aggravated its migration effects. These data suggest that the Slitrk2 function in neuronal migration relies on mechanisms distinct from those mediating synaptogenesis.

Both Slitrk1 and Slitrk2 were reported to influence neuronal morphology by affecting neuritogenesis *in vitro* [4, 35, 36]. At the final steps of migration, neurons establish their final positioning, begin to mature and develop dendrites extending into the marginal zone (MZ). We thus measured neuronal soma length and dendrites travel distance in the MZ upon OE of Slitrks and variants (**Fig. S3**). We found that neither Slitrk1 WT nor its variants altered the neuronal morphology (**Fig. S3A**). In contrast, we found significantly reduced neuronal cell bodies length and dendrites travel distance in Slitrk2 OE neurons in the upper CP (**Fig. S3B**), suggesting that neurons overexpressing Slitrk2 stopped migration, acquiring a less polarized morphology that correlates with a different distribution in the CP. Interestingly, reduced neuronal soma length was lost upon OE of both Slitrk2 variants, while their dendritic travel distance was still decreased. These results suggest that soma length and dendrites travel distance are two independent parameters and differentially influenced by the two Slitrk2 variants. In addition, as for Slitrk1, the migration phenotypes induced by OE of the Slitrk2 variants do not directly correlate with morphological changes. Together, the L627F variant enhances the cell migration effect promoted by Slitrk2 during early corticogenesis while there is no correlation between synapse organizer activity of the V89M and migration control.

In conclusion, pathological variants appear to affect Slitrk proteins function in radial migration, prior to synaptogenesis, and distinct protein domains are to be involved in the regulation of these two biological processes.

### Slitrk1 subcellular targeting is defective in truncated mutant forms

The lack of functionality in some Slitrk mutants may partially rely on their protein mislocalization, as previously showed in *in vitro* studies [4, 33, 35]. When OE *in vivo* in E13.5-born glutamatergic neurons, Slitrk1 is localized at E17.5 to the somatodendritic compartment in post-migratory neurons of the upper CP (**Fig. 3**), consistent with the endogenously distribution during cortical development [37]. Enriched expression of Slitrk1 in the dendritic compartment (magenta) is evident in the MZ, the layer composed of the apical dendrites. In both the CP and IZ stalled cells Slitrk1 accumulation was both detected in neurites and soma. Membrane protrusions were decorated by the Slitrk1 accumulation in these cells (**Fig. 3**).

**Figure 3.**
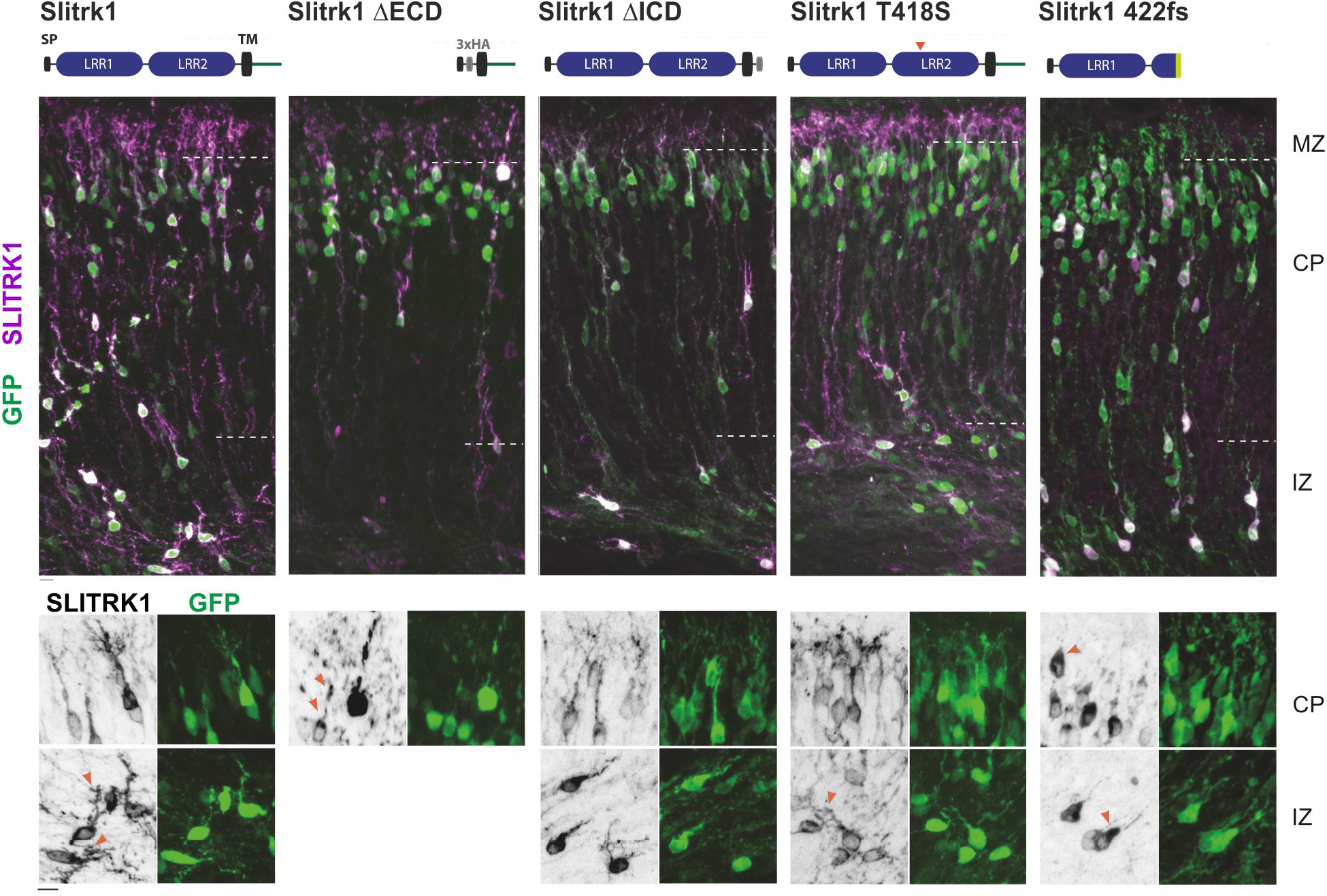
Protein localization is altered in migrating neurons overexpressing Slitrk1 variants. Immunostaining for Slitrk1 (magenta) (using either HA and/or Slitrk1 antibodies, acquired at a high detection threshold where endogenous protein is no longer visible) and GFP (green) in electroporated cells in E17.5 lateral cortex upon electroporation of Slitrk1 (WT) and its mutant forms. Slitrk1 (WT) OE is localized to the soma of postmigratory neurons in the upper CP and their dendritic compartment in the MZ (magenta top panels), and into neurites and soma in IZ stalled cells (orange arrowheads), observed in ≥ 3 embryos. In the CP, in the absence of the ECD domain (ΔECD) the protein is enriched in the soma and dendritic distribution is compromised in the MZ (orange arrowheads, n=2). The absence of ICD (ΔICD) leads to a reduced localization in dendrites: note reduced staining in the MZ and Slitrk1 puncta (black in bottom panels) in dendrites at higher magnification (n=2). Slitrk1 T418S protein distribution in CP and IZ cells is comparable to Slitrk1 (n=3). Slitrk1 422fs exhibits accumulation in the cell soma and is absent in neurites (n=3). Scale bars: 10µm.

We then compared the distribution of Slitrk1 variants. In the CP, Slitrk1 ΔECD proteins accumulated at the base of GFP^+^ dendrites and Slitrk1 ΔICD exhibited slightly altered, patchy distribution in dendrites. In the IZ, Slitrk1 ΔICD was observed in the soma and neurites of GFP^+^ cells. However, either the morphology of the cell was different from Slitrk1 OE with less ramified neurites or the mutant protein was not properly distributed. Consistently, accumulation of both the Slitrk1 ΔECD and Slitrk1 ΔICD mutant forms was not detected in the MZ. These results suggest that both Slitrk1 domains are critical for directing accurate protein localization into neurites.

Among the two disease-associated variants, Slitrk1 T418S did not manifest visible intracellular localization changes, in contrast to previously reported data, whereby surface trafficking of this mutant form was severely affected in cultures [33]. Consistently, the mutated proteins were accumulated in the MZ. Interestingly, Slitrk1 422fs (**Fig. 2A**) exhibited preferential localization to the cell soma, with most visible prevalence likely around the Golgi organelle (**Fig. 3**) in GFP^+^ cells in both the CP and IZ. This correlated with the absence of mutant Slitrk1 422fs protein accumulation in MZ dendrites. Together these observations suggest that the migration delay promoted by Slitrk1 in the CP requires a full-length protein and proper localization in dendrites/neurites and, importantly, that the Slitrk1 T418S variant behaves as the WT in both protein localization and migration.

### Cell-autonomous interaction of Slitrk1 with LAR-RPTPs occurs during discrete steps of radial migration

While Slitrk and LAR-RPTP interact in *trans*-cellular configuration, scRNAseq analyses suggested that these proteins are also co-expressed within the same cells [30] (**Fig. S4**). Moreover, Slitrk1 seemed to be co-expressed with Ptprs and Ptprd during cortical development in neurons born at E13.5 (**Fig. S4B**). Notably, Ptprs is highly expressed in young neurons (N1d, 24h after differentiation), whereas Ptprd increases in older neurons (N4d, 96h after differentiation) with Slitrk1 expressed in both. We thus examined whether Slitrks and LAR-RPTPs could cooperate in the same cell.

We first expressed Ptprs and Ptprd by IUE (**Fig. 4A-C**). Both genes induced migration delays in the CP, however different from those observed for Slitrk1 and Slitrk2. Ptprs OE cells had an overall scattered distribution in the upper cortex, with a significant decrease detected only in CP1. Ptprd OE showed even stronger scattered delay throughout the CP, reaching significance in CP1 and 3. Accumulation in the IZ was not detected for neither of them compared to controls. Next, we co-electroporated *Slitrk1* with either *Ptprs* or *Ptprd* at equal molar concentrations (**Fig. 4D, E, C**). When Slitrk1 was co-expressed with Ptprs (the MeB^-^ isoform unable to bind Slitrks ECD [6]), the delay in the upper CP, characteristic for both genes when OE separately, was abolished. These results suggest that the balanced concentrations of Slitrks and LAR-RPTPs are critical for correct termination of migration in the upper CP and that these proteins might interact in *cis* independently of the MeB. However, in co-expression experiments the Slitrk1-dependent stalled GFP^+^ population in the IZ was still present, suggesting that this phenotype is not influenced by interactions with the Ptprs isoform. On the contrary, when Slitrk1 was co-expressed with Ptprd (the MeB^+^ isoform, interacting with Slitrks LRR1 domains), we observed a reduction in GFP^+^ cells in the IZ (**Fig. 4C**) indicating that this accumulation relies on the Slitrk1 LRR1 domain and possibly the Ptprd MeB. Interestingly, Slitrk1 and Ptprd induced migration delays in the CP were differentially rescued by co-expression. The Ptprd-specific delay in CP3 were now similar to GFP controls, whereas that in CP1 characteristic of both Slitrk1 and Ptprd OE was still present. This suggests that the cell-autonomous interaction of the Slitrk1 and Ptprd proteins could be important during radial migration in the deeper rather than the upper CP. Since quantifications showed that none of the co-electroporations demonstrated purely cumulative effects (**Table S1**), direct or indirect interactions between the proteins is likely to take place. Overall, Ptprd and Ptprs seem to interact with Slitrk1 in a distinct manner during early cortical development, possibly in concordance with the temporal and spatial pattern of coexpression in young versus more mature neurons and/or in the deeper versus upper CP.

**Figure 4.**
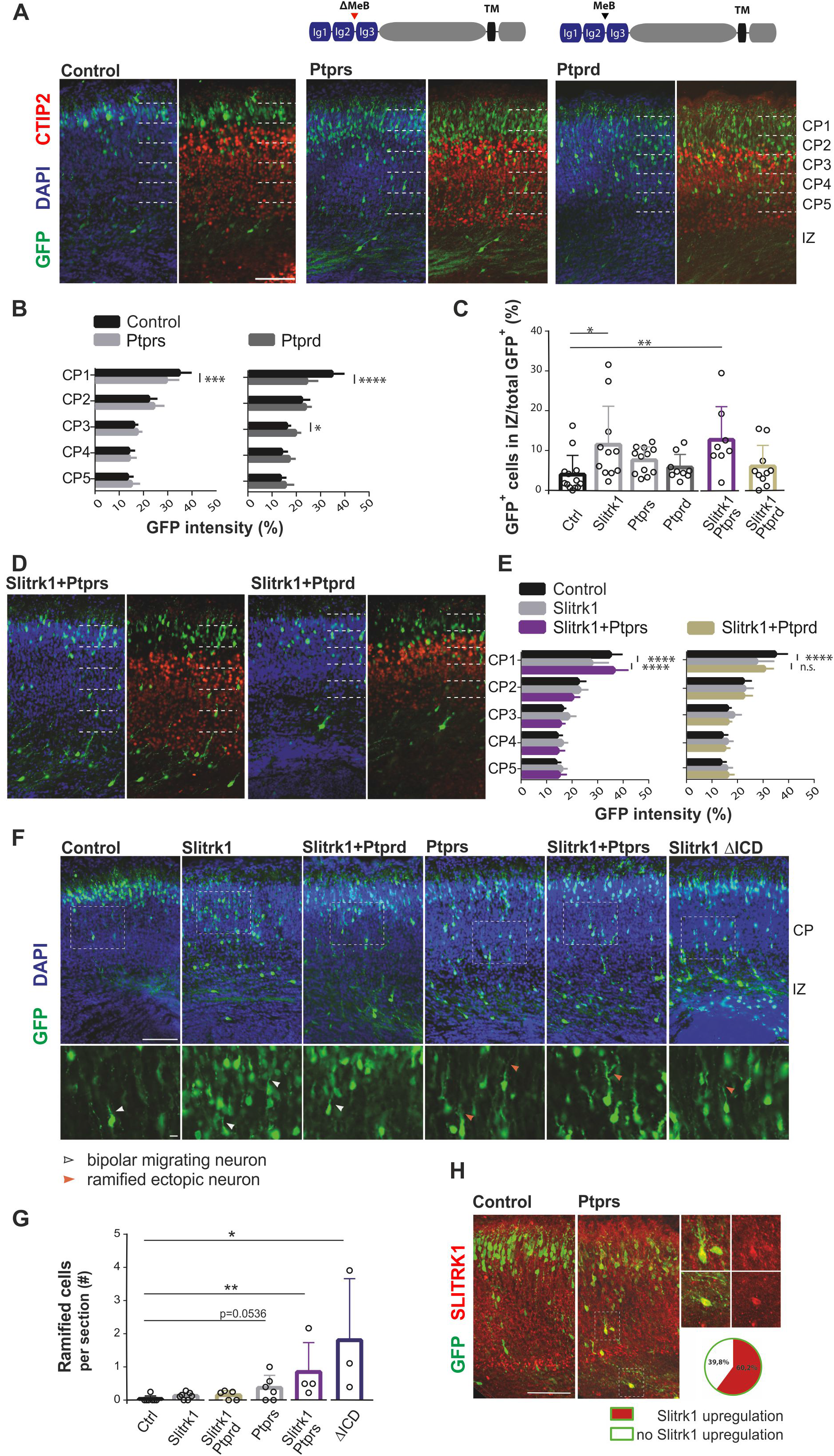
LAR-RPTPs reverse the Slitrk1 overexpression phenotype in an opposing manner and produce ectopic cells. Immunofluorescent images depicting the degrees of migration delay when Ptprs or Ptprd are overexpressed alone (**A**), and upon co-electroporation with pCAG-Slitrk1 (**D**). **B**, **E**) Quantification of migration delay in 5 equal bins, only significant differences are marked (Control *vs* Ptprs *F*(4, 105)=5.09, *P*=0.0009; *vs* Ptprd *F*(4, 90)=19.02, *P*<0.0001; Control *vs* Slitrk1 *vs* Slitrk1+Ptprs *F*(8, 150)=6.566, *P*<0.0001; Control *vs* Slitrk1 *vs* Slitrk1+Ptprd *F*(8, 135)=4.865, *P*<0.0001). **C**) Quantification of cells present in the IZ (Control *vs* Slitrk1 *vs* Ptprs *vs* Ptprd KW=10.52, *P*=0.0146; Slitrk1 *vs* Slitrk1+Ptprs *U*=36, *P*=0.5448; Control *vs* Slitrk1+Ptprd *U*=45.5, *P*=0.2374), significant differences are marked. Each point represents 1 brain section (n=8-12). **F**) Ectopic cells with enhanced ramified morphology upon overexpression of Ptprs or Ptprs with Slitrk1 and Slitrk1 ΔICD. Representative immunofluorescence images: normal bipolar migrating cells (white arrowheads) and cells ectopically developing dendrite-like arborization (orange arrowheads) are marked. **G**) Quantification of ectopic cells with arborizations per electroporated section (Control *vs* Slitrk1 *U*=12, *P*=0.0862; *vs* Slitrk1+Ptprd *U*=9.5, *P*=0.1869; *vs* Ptprs *U*=8.5, *P*=0.0536; *vs* Slitrk1+Ptprs *U*=1, *P*=0.0061; *vs* Slitrk1-ΔICD *U*=0, *P*=0.0083). Individual points represent mean value of n=3-18 sections per embryo. **H**) In pCAG-Ptprs electroporations ectopic cells accumulate Slitrk1 in 60.2% of electroporated GFP^+^ cells throughout the CP and IZ (n=3). Scale bars: 0.1mm.

In addition, we noticed a new phenotype consisting of a precocious appearance of dendrite-like ramifications in the electroporated cells before reaching the upper CP. Although detected with low frequency, these cells were specifically present in Ptprs electropored alone, Slitrk1/Ptprs co-electroporations as well as in Slitrk1 ΔICD experiments (**Fig. 4F, G**). Moreover, we detected that neurons overexpressing Ptprs upregulated endogenous Slitrk1 expression and that the majority of cells with ectopic ramifications displayed Slitrk1 upregulation (**Fig. 4H**). Together, these experiments suggest that i) LAR-RPTPs and Slitrk1 interaction may occur in *cis* independently of the MeB and/or LRR1/LRR2 domains and ii) this interaction regulates neuronal migration/arrest and maturation possibly through the ICD domain of Slitrk1.

### Slitrk1 and its variants differentially affects formation of presynaptic boutons

Since Slitrk1 overexpression promotes synaptogenesis *in vitro* [8, 10], we tested whether and how Slitrk1 OE could influence presynaptic contacts in the developing cortex. We explored mRNA expression of several presynaptic markers (**Fig. S5**) and chosed VAMP2 as its expression pattern was similar to Slitrks and LAR-RPTPs [38, 39, 40, 41], with an expression increasing with differentiation score. To assess the non-cell autonomous role of Slitrk1 on the formation of local presynaptic boutons and morphology in GFP^+^ cells we established a high-throughput protocol (**Fig. 5A**, **Movie 1, Fig. S6, Materials and Methods SI**), which automatically identifies VAMP2^+^ presynaptic boutons (VAMP2 spots) in the immediate proximity to GFP^+^ cells. We were then able to analyse VAMP2^+^ boutons properties either in “GFP^+^ dilated envelope” (noted GFP*, **Fig. 5A** in blue, **Movie 1** and **Fig. S6F**) or in the external part of those areas that are devoided of GFP^+^ cells (**Fig. S6D**).

**Figure 5.**
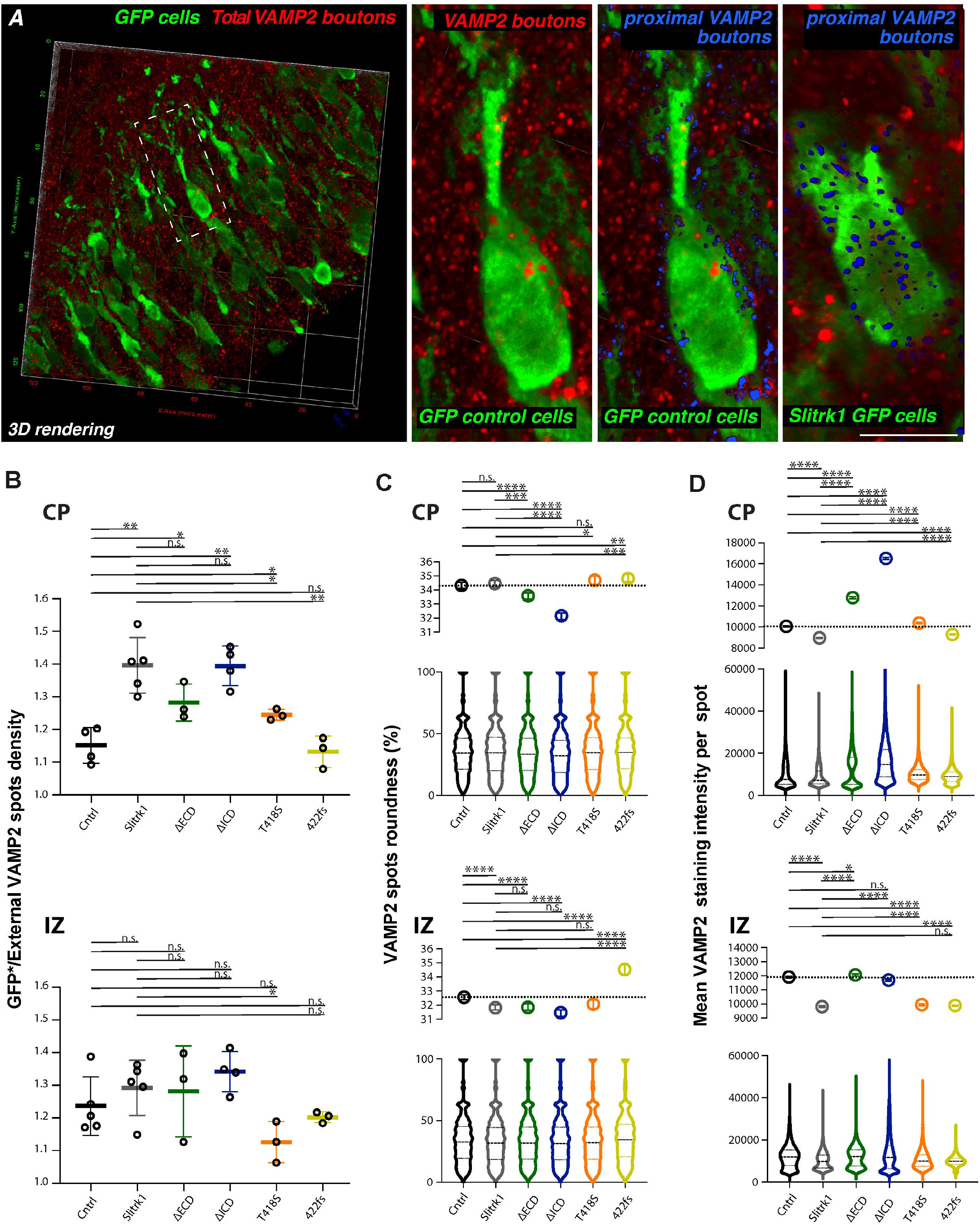
Presynaptic boutons adjacent to Slitrk1 overexpressing cells are affected in density, morphology and VAMP2 enrichment. **A**) VAMP2 spots detection in the CP by the ICY protocol. VAMP2 spots contacting either WT GFP^+^ envelope or their 3D contour (280 nm thickness) are considered as contacting boutons (in blue). GFP* cells = GFP^+^ cells + contour. Spots outside this volume are considered as external and remain in red. **B**) Quantifications of VAMP2 spots densities in GFP* cells normalized to the densities external to GFP*, CP: *F*(5, 13.82)=16, *P*<0.0001; IZ: *F*(5, 7.454)=2.776, *P*=0.1022, Brown-Forsythe ANOVA, unpaired t with Welch’s correction for multiple comparisons. Individual points represent the value per section (n=3-5). Quantifications for GFP* cells of (**C**) roundness in CP (KW=432.3, *P*<0.0001) and in IZ (KW=284.3, *P*<0.0001) and (**D**) average pixel intensity per spot in CP (KW=23010, *P*<0.0001) and in IZ (KW=6996, *P*<0.0001). Data shown as violin plots and medians with 95% CI (n=3-5).

First, we observed that in GFP* controls, VAMP2 spots densities were significantly higher in the IZ, with an overall bigger volume, higher VAMP2 staining intensities and less spherical morphology (**Fig. S7**), suggesting region-specific differences in presynaptic boutons assembly. We next examined whether presynaptic boutons could be influenced by postsynaptic overexpression of Slitrk1 and mutant forms (**Fig. 5**). In agreement with previous observations *in vitro* [8, 10] we noticed increased boutons density around the Slitrk1 OE GFP^+^ neurons in the CP (**Fig. 5B**). Interestingly, both ΔICD and ΔECD deletion mutants-expressing cells still maintained the capacity to induce high VAMP2 spots densities comparable to Slitrk1. This revealed that not only ECD, as expected, but also ICD alone, are able to promote synaptogenesis, suggesting an active role of the ICD in signaling through other *trans*-synaptic molecules. On the contrary, both disease-associated variants showed significant impairments. While T418S diminished the Slitrk1 OE effect, 422fs completely abolished it (**Fig. 5B**). Interestingly, in the IZ control and Slitrk1 OE GFP^+^ cells exhibit similar VAMP2 external spot densities. Among the variant forms, only T418S clearly showed a difference with respect to Slitrk1 OE with a significant decrease in presynaptic boutons. Since we had more pronounced effect in the CP than the IZ, this suggested that Slitrk1/partners interactions are not necessarily comparable between these regions. Moreover, since high synaptic densities did not prevent deletion mutants to correctly reach CP1, we conclude that a damaged LRR2 domain in the T418S mutant form reduces presynaptic boutons density, but not migratory behavior, of both CP and IZ neurons.

We then examined VAMP2^+^ bouton morphology by looking at roundness in 3D (**Fig. 5C**). In the CP, Slitrk1 did not affect VAMP2 synaptic shape. However, both deletion mutants, more so for ΔICD, had more elongated shape, indicating that the entire Slitrk1 is necessary to control synaptic morphology. Curiously, the effects of Slitrk1 422fs was the opposite with an increase in roundness compared to Slitrk1 OE and controls, with Slitrk1 T418S behaving as Slitrk1. This suggests that both LRR2 and ICD domains are required for proper synaptic morphology. In the IZ, both Slitrk1 and mutant forms resulted in more elongated boutons, except for the 422fs mutant that enhanced their roundness. Since this mutant form is retained in the cell (**Fig. 3**), this suggests that expression at the surface is necessary to interact with *trans*-synaptic partners and is important for bouton shape.

Lastly, we analyzed the VAMP2 fluorescence intensity (**Fig. 5D**) within boutons as this parameter is correlated with the number of synaptic vesicles. Overexpression of Slitrk1 led to a decreased VAMP2 intensity within presynaptic boutons in both the CP and the IZ. Concerning deletion mutants, we observed no difference in the IZ with respect to controls, thus preventing the Slitrk1 phenotype, but an increase in VAMP2 intensity within the CP. This correlated with the degree of spots elongation (**Table S1**). Notably, in the IZ T418S and 422fs behaved similarly to Slitrk1 with decreased VAMP2 intensity, whereas in the CP they eliminated the Slitrk1 effect. Altogether, even though Slitrk1 OE has different effects within the IZ and the CP, the overall morphology of VAMP2^+^ boutons around the neurons expressing Slitrk1 variants are strongly affected. Moreover, bouton properties in both IZ and CP do not correlate with the observed migration delays (**Table S1**), suggesting that Slitrk1-induced migration defects do not immediately depend on its capacity to neither induce presynaptic boutons assembly, nor morphology and presynaptic vesicles recruitment.

## Discussion

We show that in the embryonic cerebral cortex overexpression of Slitrks, their disease-associated mutants and LAR-RPTPs synaptic partners induces specific alterations of radial migration of glutamatergic neurons. Prior to post-natal functional synaptogenesis Slitrk1 promote VAMP2^+^ contacting boutons, whereas Slitrk1 deletion mutants are less powerful or inactive. Slitrk1 domains important for boutons recruitment differ from the ones triggering migration defects (**Table S1**). Likewise, the Slirtk1 T418S and Slitrk2 V89M disease-associated mutations, previously described to exert strong effects on synaptogenesis [33], do not alter neuronal migration, whereas the Slitrk2-L626F mutant, reported to have no effects on dendrite localization [33], promotes distinct migration delays than Slitrk2 (**Table S2**). Collectively, our results suggest novel roles for Slitrks/LAR-RPTPs in cortical development that appear dissociated from their synaptogenic activity.

### Possible effects in distinct steps of radial migration

During radial migration glutamatergic neurons change their migratory behaviors and morphology [42–44]. At the subplate (SP), multipolar migrating neurons pause, change their orientation and continue as bipolar neurons to locomote along radial fibers until they reach the upper CP edge [45]. Subsequently neurons detach from the radial glia and undergo terminal somal translocation to stop and develop dendritic arbors, which spread into the above MZ [46]. The migratory speed during glia-mediated locomotion and terminal translocation differs and is regulated through distinct molecular mechanisms [47, 48]. We showed that upregulation of Slitrk1 expression leads to two radial migration delay phenotypes. A portion of GFP^+^ neurons is accumulated in the IZ at the SP, and the other is delayed in the upper cortex, below the MZ in CP1, with a visible migration delay stretching throughout the upper half of the cortex. This suggests that Slitrk1 OE alters i) the multipolar-bipolar switch and/or pausing time below the CP, and ii) the bipolar migration speed and/or detachment from the radial glia and terminal translocation. Slitrk2 upregulating cells exhibit a delay in the upper CP, however specifically in CP2. Thus, Slitrk1 and Slitrk2 overexpression appears to alter two distinct processes occurring in the upper CP where neurons undergo soma translocation and detachment from the radial glia. In addition, we found that Slitrk2, but not Slitrk1, induces a decrease in dendrite travel distance and soma length suggesting that these two processes are important to bypass CP2 and reach CP1 and appear to occur in a Slitrk1-independent manner. Together, these results argue in favor of Slirtk1 influencing glia-mediated locomotion and likely its migration speed whereas Slirk2 rather radial glia– independent terminal translocation whereby neurons shorten their leading processes to move their cell bodies to their final position.

LAR-RPTPs OE cells exhibited scattered delay throughout the CP, with Ptprs-dependent defects accentuated in the upper CP, similarly to the Slitrk1, and Ptprd–dependent ones scattered throughout the whole CP. Both phenotypes suggest general defects in bipolar migration efficiency, however to distinct extents. Downregulation experiments were previously performed in *in vitro* assays on neurospheres/neural precursors for Ptprs/d and *in vivo* for Ptprd [49–52] and support our conclusions on their substantial role in neuronal migration.

Complementary expression patterns in young (N1d) or old neurons (N4d) ([4, 30] and **Fig. S4**) in the CP also reflect the position of migrating neurons in the IZ/lower CP or in the upper CP (1 and 4 days after birth, respectively). Endogenous Ptprd is strongly expressed in N4d and the whole CP, Ptprs in N1d and thus lower CP and IZ, Slitrk1 is enriched in SP and MZ, and Slitrk2 in VZ and lower CP. Thus, trans-interactions with cells-expressing partners during migration could specifically influence subsequential steps of migration within the IZ, lower and upper CP and support the observed delays in distinct regions of the CP.

However, we also revealed in scRNAseq data [30] co-expression of Slitrk1/2 and Ptprs/d. Consistently with experiments using the ICD domain and the opposite effects of Slitrk1 and Slitrk2 in neurite induction attributed to intracellular interactions [4, 30], we also showed that Slitrk1 and Ptprs/d influence each other activities cell-autonomously when co-electroporated. Slitrk1/Ptprs co-expression rescued migration delays in CP1, but not in the IZ, and promoted the appearance of ectopic highly ramified, resembling mature, cells. Specific rescue of the CP3 delays in co-electroporation of Ptprd and Slitrk1 indicated that delays in this region may also rely on their interactions, however not in CP1. Together with the remarkable similarity in the Ptprs OE delay pattern to that of Slitrk1 in the CP, this suggests that the balance between Slitrk1, Ptprs and Ptprd within the same cell is specifically important for several steps of bipolar migration, including migration arrest and dendrites arborization. Both the Slitrk1 ECD and ICD domains are required to influence migration indicating both extracellular and intracellular mechanisms. Intracellular interactions were shown to be important in synapse formation for Slitrk1 and Ptprd through homophilic binding [8], Slitrk2 [7, 12, 53], Ptprs [54] and their dependency on PDZ, LRR1/2 and MeB domains began to be unraveled. Our results are consistent with complex but distinct intracellular and extracellular interactions between partners and their functional domains in neuronal migration, including in differential function of Ptprd and Ptprs bearing or devoid of the MeB domain, respectively.

These data, taken together with LAR-RPTPs and Slitrks complementary expression distribution, indicate that endogenously both interactions in *cis* and *trans* could underline consecutive steps of radial migration.

### Slitrk1 mutations differentially alter subcellular localization and function

In the mouse developing cortex the Slitrk1 protein is enriched in the SP, MZ and IZ [4, 23, 37]. Interestingly, this specific protein distribution reflects its subcellular localization to the somatodendritic compartment [37] and is correctly preserved upon Slitrk1 OE. Slitrk1 ΔICD, ΔECD and 422fs mutants, however, manifest protein localization defects with the truncated Slitrk1 422fs not surprisingly being the most severely altered. These results are consistent with the glycosylation failure hypothesis for Slitrks intracellular distribution, since putative glycosylation sites are spread throughout the ECD and ICD domains [33, 35]. Their altered protein distribution along dendrites in the CPdoes not impair their synaptogenic activity, although it does affect the migration phenotype. Moreover, in contrast with previous reports of defects in dendritic targeting and synaptogenic functions in cultured neurons [33], the Slitrk1 T418S mutant behaved similarly to Slitrk1 for subcellular localization in the embryonic cortex and neuronal migration alterations. This difference could be due to cellular microenvironments (i.e. *in vivo vs in vitro*) or type of axonal inputs (thalamic vs pyramidal neurons) [55]. In addition, protein synthesis and distribution varies between neuronal types or embryos and adults [56]. Indeed, *in vitro* the Slitrk2 V89M mutation behaved differently according to cell types [33]. In our *in vivo* study, this mutant does not alter Slitrk2 effects on neuronal migration whereas the Slitrk2 L627F in the TM domain, which is correctly positioned at the membrane in dendrites [33], does. Our work thus suggests that subcellular distribution of the protein cannot solely explain the observed phenotypes and that effect in migration and synaptogenesis appears to be driven by distinct molecular mechanisms.

### Slirtks and their disease-associated variants alter neuronal migration independently of their boutons recruiting capacity

Slitrk1 overexpression leads to migration delay in the CP and triggers the recruitment of VAMP2^+^ boutons in neighboring neurons. Those contacts are less intense than in controls, indicating a potential decrease in VAMP2 vesicles. This is consistent with studies in adult CA1 pyramidal cells, whereby knockdown of Slitrk1 increases the density of docked presynaptic vesicles [55]. The truncated forms without ICD or ECD loose the capacity to alter migration, but still maintain an increased VAMP2^+^ density in the CP. Nevertheless, boutons exhibit an elongated form and higher VAMP2 intensity, with the ΔICD displaying a stronger effect, suggesting that the recruited boutons are either destabilized, or are coming from different neurons. The T418S mutation exhibits milder migration defects and bouton recruitment capability than Slitrk1 in the CP but these harbor higher VAMP2 intensity.

Slitrk1 overexpression also leads to arrested cells in the IZ that are contacted with the same density of VAMP2^+^ boutons as in control cells, however these are morphologically altered. While T418S still promoted migration arrest and boutons of similar shape, these were decreased in numbers and of larger size. These results refine our previous conclusion on the extracellular domain, showing that the LRR2 domain has a mild effect, if any, on migration and bouton shape and the LRR1 domain may be important for promoting elongated synapses. As expected, the 422fs mutation, composed of an LRR1 and a truncated LRR2 domains, which is mainly retained intracellularly, had no effect on migration and on boutons density in the CP but was still capable to alter the morphology in both the CP and the IZ suggesting that shape is controlled by intracellular interactions.

Notably, although Slitrk1 overexpression leads to migration delay in the CP and arrested cells in the IZ, the recruitment of VAMP2^+^ boutons in neighboring neurons mostly increase in the CP with little effects in the IZ. Deletion mutants alter Slitrk1 function solely in the CP. Together with the differential effects of T418S in migration and boutons assembly in the CP and the IZ, our results indicate that region-specific effect control synapse density and intensity, possibly due to the axonal tracts in IZ. Altogether our data are consistent with the work of Schroeder et al. [55], which showed that LRR proteins act in input-specific combinations and a context-dependent manner to specify synaptic properties, with LRRTM1 and Slitrk1 exerting opposite effects on synaptic vesicle docking at the active zone.

Canonical synaptogenesis mostly occurs after birth both in the mouse and primate cortex [57, 58] with a marked increase between P4 and P32. We show here in the embryonic cortex that Slitrk1 expression promotes the recruitment of VAMP2^+^ boutons over GFP^+^ cells, suggesting the formation of transient contacts with neighboring neurons. Transient contacts were shown in the hippocampus whereby distinct pioneer neurons are involved in the guidance and targeting of different hippocampal afferents [59, 60]. VAMP2 is expressed both at inhibitory and excitatory synapse so we could not infer the nature of those contacts. However, we show that morphology of VAMP2 recruited spots varies depending on the Slitrk1 protein domains.

Indeed, Slitrk1 ΔICD recruited more elongated boutons in the CP compared to Slitrk1, which are characteristic of inhibitory contacts as shown in hippocampal neurons [61]. This suggests that Slitrk1 could possibly recruit both excitatory and inhibitory transient contacts, but that mutant forms (like ΔICD) are losing excitatory contact recruitments. An alternative hypothesis, would be that remaining recruited boutons are destabilized by an altered trans-synaptic contact (with ΔECD for example), leading to a compensatory increase of synaptic vesicle content to the remaining contact. It remains to be determined whether Slitrk/LAR-RPTP OE stimulate electrical activity in migrating neurons through these contacts in the CP even before fully functional synapses are formed [43].

Altogether, our data suggests that the control of migration as well as that of density, morphology and intensity of presynaptic boutons rely on distinct molecular mechanisms involving both *trans*- and *cis*-interactions.

### Slirtks and LAR-RPTPs variants and psychiatric disorders beyond synaptopathies

Slirtks and LAR-RPTPs variants are associated with multiple psychiatric disorders, ranging from OCD and Gilles de la Tourette’s syndrome to ASD and SCZ, addictive behaviors and cognitive impairments [2, 3, 14–18]. However, all proteins are vastly expressed in the nervous system from E11 prior to functional synapse formation [4, 30]. If Slitrk1 and Slirtk2 were long known to be involved in neuritogenesis with opposite activities *in vitro* [4, 11, 19, 20], their function was mostly studied in synapse organization and activity in cultured hippocampal neurons whereby they play a crucial *trans*-synaptic role. In the last few years, several studies began to unravel their specific *cis*-activities in synaptogenesis in postnatal hippocampal development in mice and in cultured neurons. Recently, also knock-out mice in the postnatal hippocampus shed light on their differential role in the control of synapse density and electrical activity [52, 62]. Our work provides solid evidence for their role in embryonic neuronal migration in addition to the functional characterization of missense variants associated with NDDs in this process. This opens new venues on the understanding of the relevance of these mutations in NDDs extending to a novel function that was so far unknown in embryonic neocortex development. Migration, positioning and maturation of neurons precedes and is a prerequisite for the establishment of functional neuronal circuits. We show that mutations exert distinct effects in neuronal migration and in recruiting VAMP2^+^ boutons with surprisingly uncorrelated functions in these two processes. Altogether, our data strongly argue in favor of Slirtks and LAR-RPTPs-associated pathologies being the results of earlier embryonic defects in neuronal migration in addition to pure “canonical” synaptopathies.

## Supporting information

Medvedeva Supplemntary Figures

## Acknowledgments

VPM was supported by the Fondation Imagine (ANR-10-IAHU-01) and AP is a CNRS (Centre National de la Recherche Scientifique) Investigator. PB and LD are INSERM (Institut National de Santé et Recherche Médicale) researchers. This work was supported by grants from the Agence Nationale de la Recherche (ANR-15-CE16-0003-01), FRM («Equipe FRM DEQ20130326521»), and Fondation de France (00081243) to AP and FLAG-ERA grant Senseï by ANR-19-HBPR-0003 to LD.

We would like to thank Quentin Dholandre (Institute Imagine) for technical assistance with cryosectioning, the Neurimag Imaging Facility team (part of IPNP, Inserm U. 1266 and Université Paris Cité) for their technical and scientific support in imaging of biological sample. We thank the Leducq foundation for supporting the acquisition of the Leica SP8 Confocal/STED 3DX microscope.

## Author contributions

Conceptualization: VM, PB, LD and AP

Methodology: VM, PB, AJ, LV, JK, LD and AP

Investigation: VM, PB, LD and AP

Data Curation: VM, PB, LD and AP

Writing – original draft preparation: VM, PB, LD and AP

Writing – review and editing: VM, PB, AJ, LV, JK, LD and AP

Visualization: VM, PB, LD and AP

Contacting synapse software: LD

Supervision: AP

Project administration: AP

Funding acquisition: LD, AP

## References

1. Ko J. The leucine-rich repeat superfamily of synaptic adhesion molecules: LRRTMs and slitrks. Molecules and Cells. 2012;34:335–340.

2. Takahashi H, Craig AM. Protein tyrosine phosphatases PTPδ, PTPς, and LAR: Presynaptic hubs for synapse organization. Trends in Neurosciences. 2013;36:522–534.

3. Um JW, Ko J. LAR-RPTPs: Synaptic adhesion molecules that shape synapse development. Trends in Cell Biology. 2013;23:465–475.

4. Aruga J, Mikoshiba K. Identification and characterization of Slitrk, a novel neuronal transmembrane protein family controlling neurite outgrowth. Molecular and Cellular Neuroscience. 2003;24:117–129.

5. Chagnon MJ, Uetani N, Tremblay ML. Functional significance of the LAR receptor protein tyrosine phosphatase family in development and diseases. Biochem Cell Biol. 2004;82:664–675.

6. Um JW, Kim KH, Park BS, Choi Y, Kim D, Kim CY, et al. Structural basis for LAR-RPTP/Slitrk complex-mediated synaptic adhesion. Nature Communications. 2014;5:5423.

7. Yamagata A, Sato Y, Goto-Ito S, Uemura T, Maeda A, Shiroshima T, et al. Structure of Slitrk2-PTPδ complex reveals mechanisms for splicing-dependent trans-synaptic adhesion. Scientific Reports. 2015;5:9686.

8. Beaubien F, Raja R, Kennedy TE, Fournier AE, Cloutier JF. Slitrk1 is localized to excitatory synapses and promotes their development. Scientific Reports. 2016;6:27343.

9. Won SY, Lee P, Kim HM. Synaptic organizer: Slitrks and type IIa receptor protein tyrosine phosphatasess. Current Opinion in Structural Biology. 2019;54:95–103.

10. Yim YS, Kwon Y, Nam J, Yoon HI, Lee K, Kim DG, et al. Slitrks control excitatory and inhibitory synapse formation with LAR receptor protein tyrosine phosphatases. Proceedings of the National Academy of Sciences. 2013;110:4057–4062.

11. Salesse C, Charest J, Doucet-Beaupré H, Castonguay AM, Labrecque S, De Koninck P, et al. Opposite Control of Excitatory and Inhibitory Synapse Formation by Slitrk2 and Slitrk5 on Dopamine Neurons Modulates Hyperactivity Behavior. Cell Reports. 2020;30:2374–2386.e5.

12. Han KA, Kim J, Kim H, Kim D, Lim D, Ko J, et al. Slitrk2 controls excitatory synapse development via PDZ-mediated protein interactions. Scientific Reports. 2019;9.

13. Li J, Han W, Pelkey KA, Duan J, Mao X, Wang YX, et al. Molecular Dissection of Neuroligin 2 and Slitrk3 Reveals an Essential Framework for GABAergic Synapse Development. Neuron. 2017;96:808–826.e8.

14. Piton A, Gauthier J, Hamdan FF, Lafrenière RG, Yang Y, Henrion E, et al. Systematic resequencing of X-chromosome synaptic genes in autism spectrum disorder and schizophrenia. Molecular Psychiatry. 2011;16:867–880.

15. Proenca CC, Gao KP, Shmelkov S V, Rafii S, Lee FS. Slitrks as emerging candidate genes involved in neuropsychiatric disorders. Trends in Neurosciences. 2011;34:143–153.

16. Bansal V, Mitjans M, Burik CAP, Linnér RK, Okbay A, Rietveld CA, et al. Genome-wide association study results for educational attainment aid in identifying genetic heterogeneity of schizophrenia. Nature Communications. 2018;9.

17. Depienne C, Ciura S, Trouillard O, Bouteiller D, Leitã O E, Nava C, et al. Association of Rare Genetic Variants in Opioid Receptors with Tourette Syndrome. Tremor and Other Hyperkinetic Movements (New York, NY). 2019;9.

18. Uhl GR, Martinez MJ. PTPRD: neurobiology, genetics, and initial pharmacology of a pleiotropic contributor to brain phenotypes. Annals of the New York Academy of Sciences. 2019;1451:112–129.

19. Wang J, Bixby JL. Receptor tyrosine phosphatase-δ is a homophilic, neurite-promoting cell adhesion molecule for CNS neurons. Molecular and Cellular Neurosciences. 1999;14:370–384.

20. Thompson KM, Uetani N, Manitt C, Elchebly M, Tremblay ML, Kennedy TE. Receptor protein tyrosine phosphatase sigma inhibits axonal regeneration and the rate of axon extension. Molecular and Cellular Neuroscience. 2003;23:681–692.

21. Sommer L, Rao M, Anderson DJ. RPTPδ and the novel protein tyrosine phosphatase RPTPψ are expressed in restricted regions of the developing central nervous system. Developmental Dynamics. 1997;208:48–61.

22. Schaapveld RQ, Schepens JT, Bächner D, Attema J, Wieringa B, Jap PH, et al. Developmental expression of the cell adhesion molecule-like protein tyrosine phosphatases LAR, RPTPdelta and RPTPsigma in the mouse. Mechanisms of Development. 1998;77:59– 62.

23. Beaubien F, Cloutier JF. Differential expression of Slitrk family members in the mouse nervous system. Developmental Dynamics. 2009;238:3285–3296.

24. Saito T. In vivo electroporation in the embryonic mouse central nervous system. Nature Protocols. 2006;1:1552–1558.

25. Matsui A, Yoshida AC, Kubota M, Ogawa M, Shimogori T. Mouse in utero electroporation: Controlled spatiotemporal gene transfection. Journal of Visualized Experiments. 2011:1–5.

26. Barber M, Arai Y, Morishita Y, Vigier L, Causeret F, Borello U, et al. Migration speed of Cajal-Retzius cells modulated by vesicular trafficking controls the size of higher-order cortical areas. Current Biology. 2015;25:2466–2478.

27. De Chaumont F, Dallongeville S, Chenouard N, Hervé N, Pop S, Provoost T, et al. Icy: An open bioimage informatics platform for extended reproducible research. Nature Methods. 2012;9:690–696.

28. Lagache T, Grassart A, Dallongeville S, Faklaris O, Sauvonnet N, Dufour A, et al. Mapping molecular assemblies with fluorescence microscopy and object-based spatial statistics. Nature Communications 2018 9:1. 2018;9:1–15.

29. Anton-Sanchez L, Bielza C, Merchán-Pérez A, Rodríguez JR, DeFelipe J, Larrañaga P. Three-dimensional distribution of cortical synapses: A replicated point pattern-based analysis. Frontiers in Neuroanatomy. 2014;8:85.

30. Telley L, Agirman G, Prados J, Amberg N, Fièvre S, Oberst P, et al. Temporal patterning of apical progenitors and their daughter neurons in the developing neocortex. Science (New York, NY). 2019;364.

31. Ozomaro U, Cai G, Kajiwara Y, Yoon S, Makarov V, Delorme R, et al. Characterization of SLITRK1 Variation in Obsessive-Compulsive Disorder. PLoS ONE. 2013;8:e70376.

32. Alexander J, Potamianou H, Xing J, Deng L, Karagiannidis I, Tsetsos F, et al. Targeted Re-sequencing approach of candidate genes implicates rare potentially functional variants in tourette syndrome etiology. Frontiers in Neuroscience. 2016;10.

33. Kang H, Han K, Won SY, Kim HM, Lee YH, Ko J, et al. Slitrk missense mutations associated with neuropsychiatric disorders distinctively impair slitrk trafficking and synapse formation. Frontiers in Molecular Neuroscience. 2016;9:1–18.

34. Abelson JF, Kwan KY, O’Roak BJ, Baek DY, Stillman AA, Morgan TM, et al. Sequence variants in SLITRK1 are associated with Tourette’s syndrome. Science. 2005;310:317–320.

35. Kajiwara Y, Buxbaum JD, Grice DE. SLITRK1 Binds 14-3-3 and Regulates Neurite Outgrowth in a Phosphorylation-Dependent Manner. Biological Psychiatry. 2009;66:918– 925.

36. Sareen D, O’Rourke JG, Meera P, Muhammad AKMG, Grant S, Simpkinson M, et al. Targeting RNA foci in iPSC-derived motor neurons from ALS patients with a C9ORF72 repeat expansion. Science Translational Medicine. 2013;5:208ra149.

37. Stillman AA, Krsnik Ž, Sun J, Rašin MR, State MW, Şestan N, et al. Developmentally regulated and evolutionarily conserved expression of SLITRK1 in brain circuits implicated in Tourette syndrome. Journal of Comparative Neurology. 2009;513:21–37.

38. Chen YA, Scheller RH. SNARE-mediated membrane fusion. Nature Reviews Molecular Cell Biology 2001 2:2. 2001;2:98–106.

39. Schoch S, Deák F, Königstorfer A, Mozhayeva M, Sara Y, Südhof TC, et al. SNARE function analyzed in synaptobrevin/VAMP knockout mice. Science. 2001;294:1117–1122.

40. Ahmari SE, Buchanan JA, Smith SJ. Assembly of presynaptic active zones from cytoplasmic transport packets. Nature Neuroscience. 2000;3:445–451.

41. Zhai R, Olias G, Chung WJ, Lester RAJ, Tom Dieck S, Langnaese K, et al. Temporal appearance of the presynaptic cytomatrix protein bassoon during synaptogenesis. Molecular and Cellular Neurosciences. 2000;15:417–428.

42. Ohtaka-Maruyama C, Okado H. Molecular pathways underlying projection neuron production and migration during cerebral cortical development. Frontiers in Neuroscience. 2015;9:447.

43. Medvedeva VP, Pierani A. How Do Electric Fields Coordinate Neuronal Migration and Maturation in the Developing Cortex? Frontiers in Cell and Developmental Biology. 2020;8:580657.

44. Ohtaka-Maruyama C. Subplate Neurons as an Organizer of Mammalian Neocortical Development. Frontiers in Neuroanatomy. 2020;14:8.

45. Hirota Y, Nakajima K. Control of Neuronal Migration and Aggregation by Reelin Signaling in the Developing Cerebral Cortex. Frontiers in Cell and Developmental Biology. 2017;5:40.

46. Hirota Y, Nakajima K. Control of Neuronal Migration and Aggregation by Reelin Signaling in the Developing Cerebral Cortex. Frontiers in Cell and Developmental Biology. 2017;5:40.

47. Heck N, Kilb W, Reiprich P, Kubota H, Furukawa T, Fukuda A, et al. GABA-A receptors regulate neocortical neuronal migration in vitro and in vivo. Cerebral Cortex. 2007;17:138–148.

48. Hurni N, Kolodziejczak M, Tomasello U, Badia J, Jacobshagen M, Prados J, et al. Transient Cell-intrinsic Activity Regulates the Migration and Laminar Positioning of Cortical Projection Neurons. Cerebral Cortex. 2017;27:3052–3063.

49. Meathrel K, Adamek T, Batt J, Rotin D, Doering LC. Protein tyrosine phosphatase σ-deficient mice show aberrant cytoarchitecture and structural abnormalities in the central nervous system. Journal of Neuroscience Research. 2002;70:24–35.

50. Kirkham DL, Pacey LKK, Axford MM, Siu R, Rotin D, Doering LC. Neural stem cells from protein tyrosine phosphatase sigma knockout mice generate an altered neuronal phenotype in culture. BMC Neuroscience. 2006;7:50.

51. Zhong J, Lan C, Zhang C, Yang Y, Chen WX, Zhang KY, et al. Chondroitin sulfate proteoglycan represses neural stem/progenitor cells migration via PTPσ/α-actinin4 signaling pathway. Journal of Cellular Biochemistry. 2019;120:11008–11021.

52. Tomita H, Cornejo F, Aranda-Pino B, Woodard CL, Rioseco CC, Neel BG, et al. The Protein Tyrosine Phosphatase Receptor Delta Regulates Developmental Neurogenesis. Cell Reports. 2020;30:215–228.e5.

53. Loomis C, Stephens A, Janicot R, Baqai U, Drebushenko L, Round J. Identification of MAGUK scaffold proteins as intracellular binding partners of synaptic adhesion protein Slitrk2. Molecular and Cellular Neuroscience. 2020;103.

54. Won SY, Kim CY, Kim D, Ko J, Um JW, Lee SB, et al. LAR-rptp clustering is modulated by competitive binding between synaptic adhesion partners and heparan sulfate. Frontiers in Molecular Neuroscience. 2017;10.

55. Schroeder A, Vanderlinden J, Vints K, Ribeiro LF, Vennekens KM, Gounko N V., et al. A Modular Organization of LRR Protein-Mediated Synaptic Adhesion Defines Synapse Identity. Neuron. 2018;99:329–344.e7.

56. Mofatteh M. mRNA localization and local translation in neurons. AIMS Neuroscience. 2020;7:299–310.

57. Bourgeois JP, Rakic P. Changes of synaptic density in the primary visual cortex of the macaque monkey from fetal to adult stage. The Journal of Neuroscience : The Official Journal of the Society for Neuroscience. 1993;13:2801–2820.

58. De Felipe J, Marco P, Fairén A, Jones EG. Inhibitory synaptogenesis in mouse somatosensory cortex. Cerebral Cortex (New York, NY : 1991). 1997;7:619–634.

59. Supèr H, Martínez A, Del Río JA, Soriano E. Involvement of distinct pioneer neurons in the formation of layer-specific connections in the hippocampus. The Journal of Neuroscience : The Official Journal of the Society for Neuroscience. 1998;18:4616–4626.

60. Supèr H, Soriano E, Uylings HBM. The functions of the preplate in development and evolution of the neocortex and hippocampus. Brain Research Reviews. 1998;27:40–64.

61. Danglot L, Rostaing P, Triller A, Bessis A. Morphologically identified glycinergic synapses in the hippocampus. Molecular and Cellular Neuroscience. 2004;27:394–403.

62. Sclip A, Südhof TC. LAR receptor phospho-tyrosine phosphatases regulate NMDA-receptor responses. ELife. 2020;9.

63. Züchner S, Cuccaro ML, Tran-Viet KN, Cope H, Krishnan RR, Pericak-Vance MA, et al. SLITRK1 mutations in trichotillomania. Molecular Psychiatry. 2006;11:888–889.

64. Miranda DM, Wigg K, Kabia EM, Feng Y, Sandor P, Barr CL. Association of SLITRK1 to Gilles de la Tourette Syndrome. American Journal of Medical Genetics, Part B: Neuropsychiatric Genetics. 2009;150:483–486.

