## Supplementary material for "Slitrk/LAR-RPTP and disease-associated variants control neuronal migration in the developing mouse cortex independently of synaptic organizer activity": Medvedeva Supplemntary Figures

### Supplemental Figures

#### Supplementary Figure Legends

**Figure S1. Validation of Slitrks overexpression.** *In situ* hybridation with Slitrk1, Slitrk2 and Slitrk3 mRNAs (violet) on E17.5 neocortical sections from control and electroporated embryos for each gene confirming overexpression. Immunohistochemistry for GFP and counterstaining with DAPI were used to detect the electroporated areas and to delineate the CP ad the IZ. Scale bar: 0.1mm.

**Figure S2. pCAG-Slitrk1 delayed cells do not change their identity in the IZ or CP.** Representative confocal images in controls and Slitrk1 OE at E17.5 of immunofluorescence using: **A**) GFP (green), BRN2 (red) and CTIP2 (blue) showing that most $GFP^{+}$ cells co-express BRN2 as expected from IUE at E13.5; **B**) GFP (green) and PH3 or PAX6 or TBR2 or activated-Caspase 3 (magenta) showing that electroporated cells are not expressing markers of apical and basal progenitors or are undergoing cell death; **C**) GFP (green) and NURR1 (NR4A2) or CTGF (magenta) showing that electroporated cells are not expressing subplate (SP) markers and are positioned just below the SP or in the CP; **D**) GFP (green) and TUJ1 or TBR1 or GABA (magenta) showing that electroporated cells are neurons not expressing markers of layer 6 or of GABAergic neurons. Overall GFP intensity in the CP was comparable to control (t(6)=0.5291, P= 0.6158, unpaired t-test)).

**Figure S3**. **Slitrk1 and Slitrk2 overexpression and their NDDs-associated variants have distinct effects on morphology of cortical neurons.** **A**) Representative images of electroporations of Slitrk1 and its mutant forms in upper CP neurons of the lateral cortex. Above: schema depicting the measuring principle of the dendrite travel distance in the MZ and soma length (wider diameter). Below: an example of cell morphology found in the IZ upon Slitrk1 OE, and quantification results, where no significant differences were detected. Points represent individual dendrite or cell measurements (n=3-4). **B**) Representative images of electroporations of Slitrk2 and its variants in upper CP neurons of the lateral cortex. Below: quantifications, only significant differences are marked: soma length (Control *vs* Slitrk2 t(79)=2.616, *P*=0.0106; Slitrk2 *vs* Slitrk2-V89M t(85)=3.468, *P*=0.0008; Slitrk2 *vs* Slitrk2 L627F t(54)=2.839, *P*=0.0064; unpaired t-tests) and dendrite travel distance (Control *vs* Slitrk2 t(36)=2.408, *P*=0.0213; Slitrk2 *vs* Slitrk2 V89M t(36)=0.4586, *P*=0.6493; Slitrk2 *vs* Slitrk2 L627F t(37)=0.1087, *P*=0.9140; unpaired t-tests). Points represent individual dendrite or cell measurements (n=3-4). Scale bars=10µm.

**Figure S4. Slitrk1, Ptprs and Ptprd co-expression in early cortical development. A**) Data from Telley et al. (Telley et al., 2019). A high resolution method named Flash tag (FT) was used to label apical progenitors and their daughter neurons to trace their transcription trajectory from E12 to E15 (when layer L6, L5, L4 and L2/3 are successively generated). 1h, 24h or 96h after labeling, apical progenitors that were positive for FT were microdissected from the somatosensory cortex, sorted by FACS and submitted to single-cell RNA sequencing (scRNAseq). These data were generated on 4 lineages of cortical progenitors. We extracted the expression dynamics of the four genes of interest for the present paper (Slitrk1, Slitrk2, Ptprd and Ptprs) from the genebrowser database associated to the paper (http://genebrowser.unige.ch/telagirdon/#query_the_atlas). Note that the population investigated in our study (purple rectangle) is corresponding to the E13-E17 lineage. **B**) Stochastic neighbor embedding (t-SNE) was used to analyse cellular transcriptional identities (Telley et al., 2019). We extracted gene expression on t-SNE representation of the scRNAseq dataset specifically for Slitrk1, Slitrk2, Ptprd and Ptprs. E13-E17 neurons (N4d) are zoomed up. Orange arrowheads: N4d neurons co-expressing Slitrk1 and RPTPs. AP, apical progenitors; Astro, astrocytes; BP, basal progenitors; IN, interneurons; N1d, 1-day-old neurons; N4d, 4-day-old neurons.

**Figure S5. scRNAseq of other presynaptic proteins from Telley et al., 2019.** mRNA expression of several presynaptic markers from E12 to E15 (vesicular SNARES (vSNAREs) (VAMP1, 2, 3, 4 and 7), or dedicated glutamatergic (Slc17a6, a7 and a8) and GABAergic (GAD1 and Slc32a1) markers) showing their distinct temporal expression profiles during progenitor to neuron maturation. VAMP2 has the closest expression pattern to those of Slitrks and PTPRs, with an expression increasing with differentiation score.

**Figure S6**. **Presynaptic boutons detection strategy by the ICY protocol, as shown on CP1 control cells**. To estimate the region of synaptic junction (VAMP2 staining in **B**) touching $GFP^+$ (**A**) cells, we segmented the $GFP^+$ cells in 3D (**C**) and slightly dilate the 3D envelope of a few hundreds nanometers to englobe touching presynaptic boutons (**E**). We then analysed the VAMP2 presynaptic boutons density and shape either in "$GFP^+$ dilated envelope" (noted $GFP^*$) (**F**), also **Fig. 7A** in blue, **Movie 1**, or in the external part of those areas that are devoid of $GFP^+$ cells (**D**).

**Figure S7. Presynaptic boutons differences in the upper CP and IZ in GFP* controls. A**) VAMP2 spots density in GFP* cells is higher in IZ (t(3)=8.123, *P*=0.0039; paired t-test). This is similar to VAMP2 density data external to GFP*: mean IZ/CP spots density=1.374, t(3)=4.863, *P*=0.0166; paired t-test. Individual points represent the value per section. **B**) VAMP2 spots roundness is overall lower in IZ (*U*=1745443541, *P*<0.0001), data shown as violin plots and medians with 95% CI (n=4-5). **C**) VAMP2 spots volume is overall higher in IZ (left graph: violin plots and geometric means with 95% CI, *U*=1795754055, *P*<0.0001), right graph - spots volume minimal binning highlights 3 size groups (approximate diameter is indicated) which differ between the areas (*F*(6, 42)=2.978, *P*=0.0163, two-way ANOVA, uncorrected Fisher's LSD for multiple comparisons), data shown as mean with SEM. **D**) VAMP2 average pixel intensity is lower in the upper CP (*U*=1327263032, *P*<0.0001), data shown as violin plots and medians with 95% CI (n=4-5).

**Table S1**. A summary of migration and synaptogenic phenotypes described for Slitrk1, its mutant forms and RPTPs OE in E13.5-17.5 electroporations.

**Table S2.** Slitrk2 protein domains and disease-associated variants in radial migration.

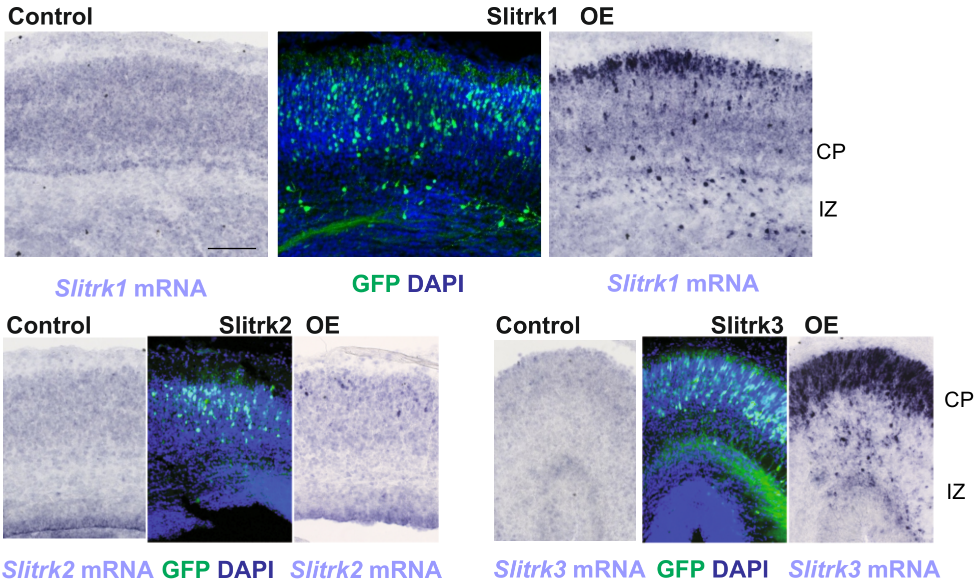
Control
Slitrk1 OE
CP
IZ
*Slitrk1* mRNA
GFP DAPI
*Slitrk1* mRNA
Control
Slitrk2 OE
Control
Slitrk3 OE
CP
IZ
*Slitrk2* mRNA
GFP DAPI
*Slitrk2* mRNA
*Slitrk3* mRNA
GFP DAPI
*Slitrk3* mRNA

### Figure S1

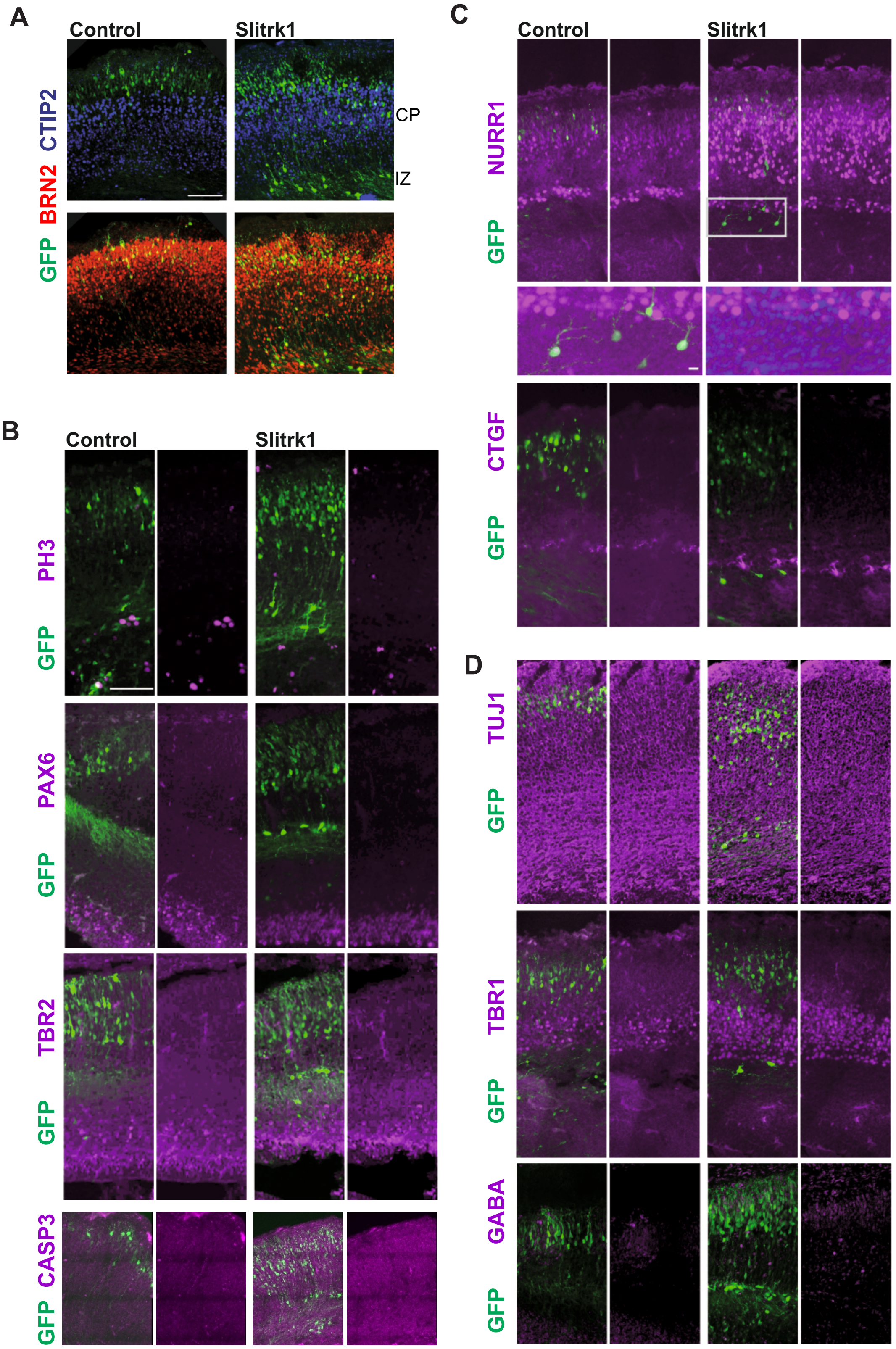
A
Control
Slitrk1
CTIP2
BRN2
GFP
CP
IZ
B
Control
Slitrk1
PH3
GFP
PAX6
GFP
TBR2
GFP
CASP3
GFP
C
Control
Slitrk1
NURR1
GFP
CTGF
GFP
D
TUJ1
GFP
TBR1
GFP
GABA
GFP

**Figure S2**

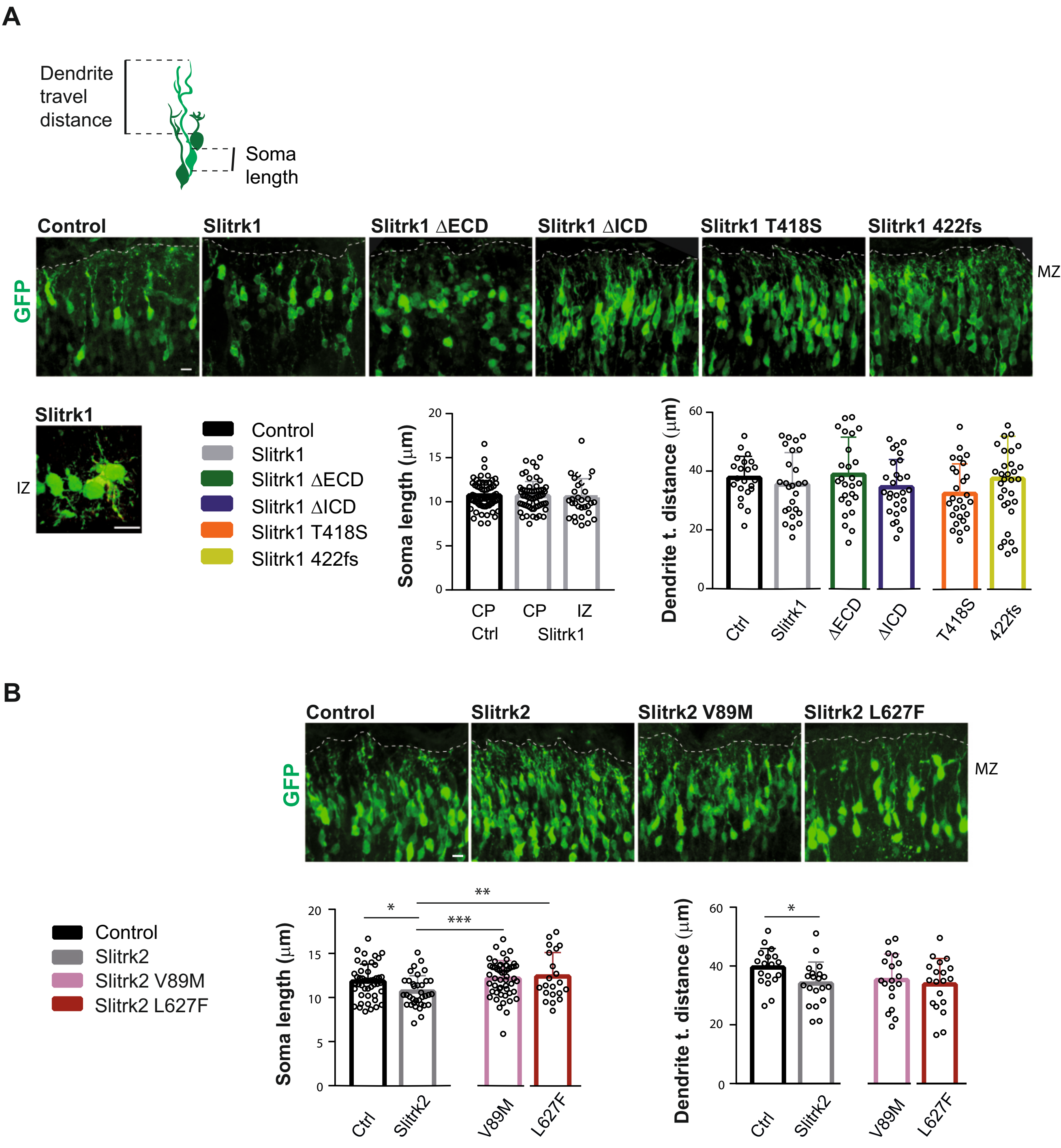

A
Dendrite travel distance
Soma length
Control
Slitrk1
Slitrk1 ΔECD
Slitrk1 ΔICD
Slitrk1 T418S
Slitrk1 422fs
GFP
MZ
Slitrk1
IZ
Control
Slitrk1
Slitrk1 ΔECD
Slitrk1 ΔICD
Slitrk1 T418S
Slitrk1 422fs
Soma length (μm)
CP
CP
IZ
Ctrl
Slitrk1
Dendrite t. distance (μm)
Ctrl
Slitrk1
ΔECD
ΔICD
T418S
422fs
B
Control
Slitrk2
Slitrk2 V89M
Slitrk2 L627F
GFP
MZ
Control
Slitrk2
Slitrk2 V89M
Slitrk2 L627F
Soma length (μm)
Ctrl
Slitrk2
V89M
L627F
Dendrite t. distance (μm)
Ctrl
Slitrk2
V89M
L627F


Figure S3

**A**

**Temporal patterning in the developing cortex**
Telley et al., Science 2019

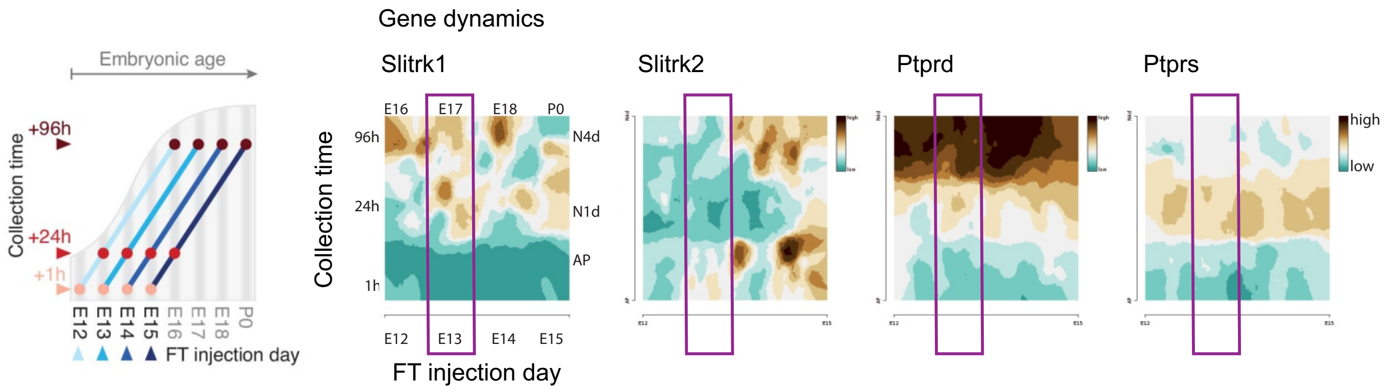


**B**

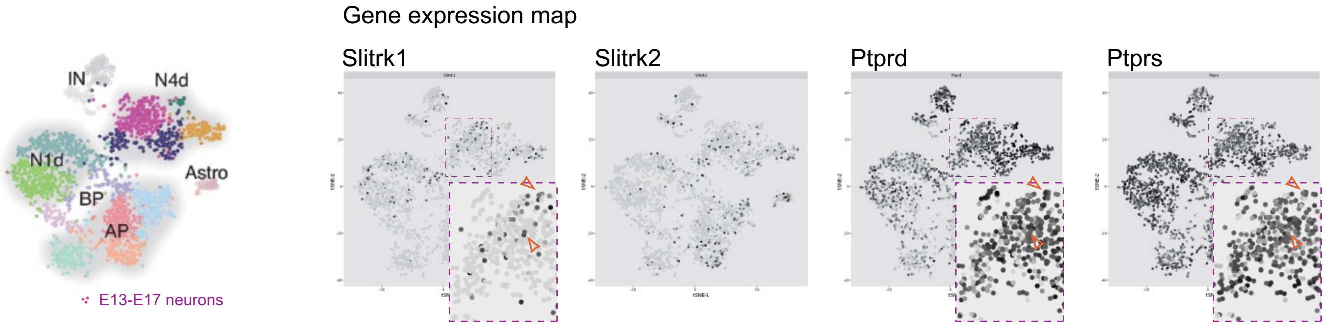


### Figure S4

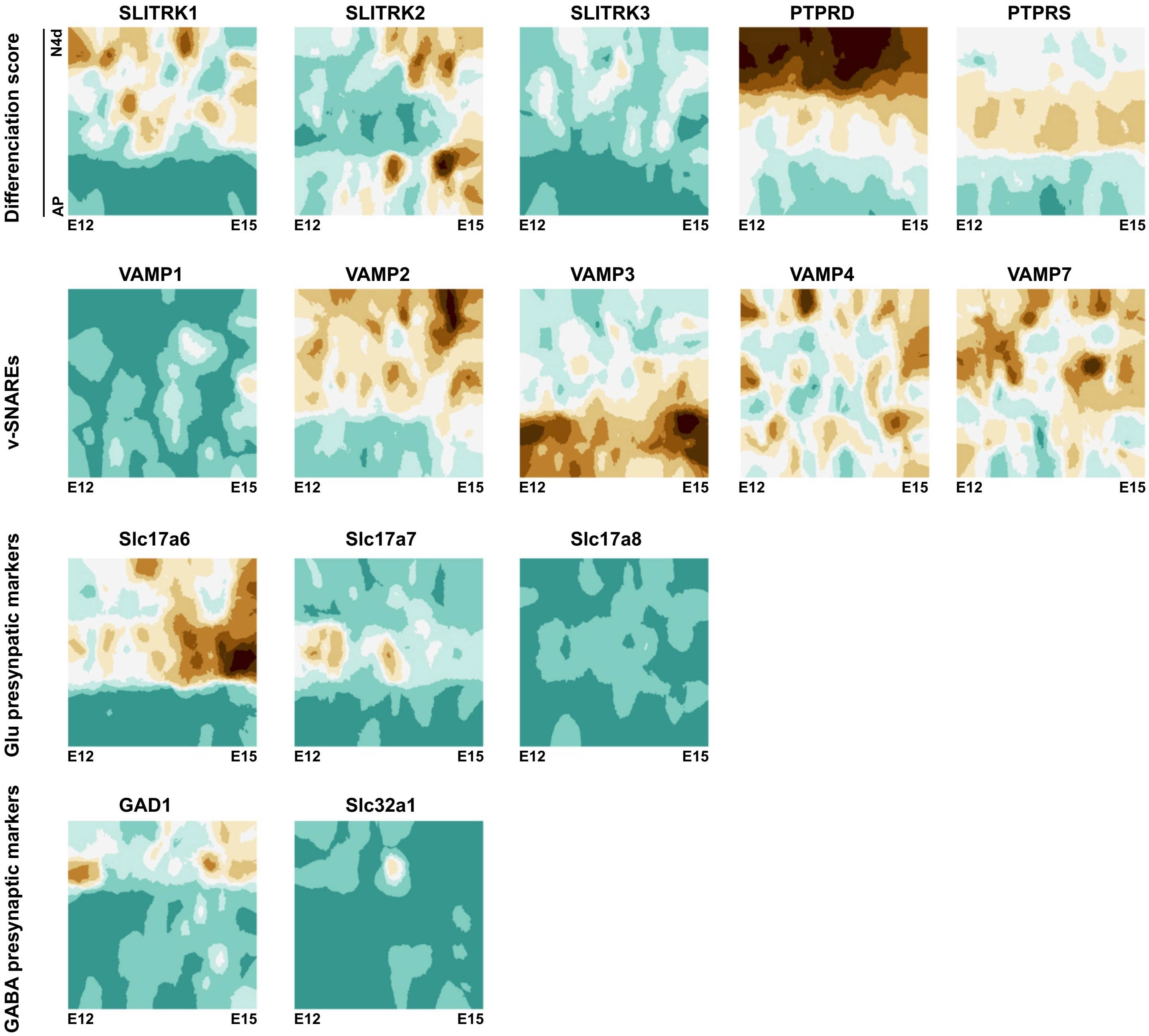

Differenciation score
N4d
AP
SLITRK1
SLITRK2
SLITRK3
PTPRD
PTPRS
v-SNAREs
VAMP1
VAMP2
VAMP3
VAMP4
VAMP7
Glu presynpatic markers
Slc17a6
Slc17a7
Slc17a8
GABA presynaptic markers
GAD1
Slc32a1
E12
E15


### Figure S5

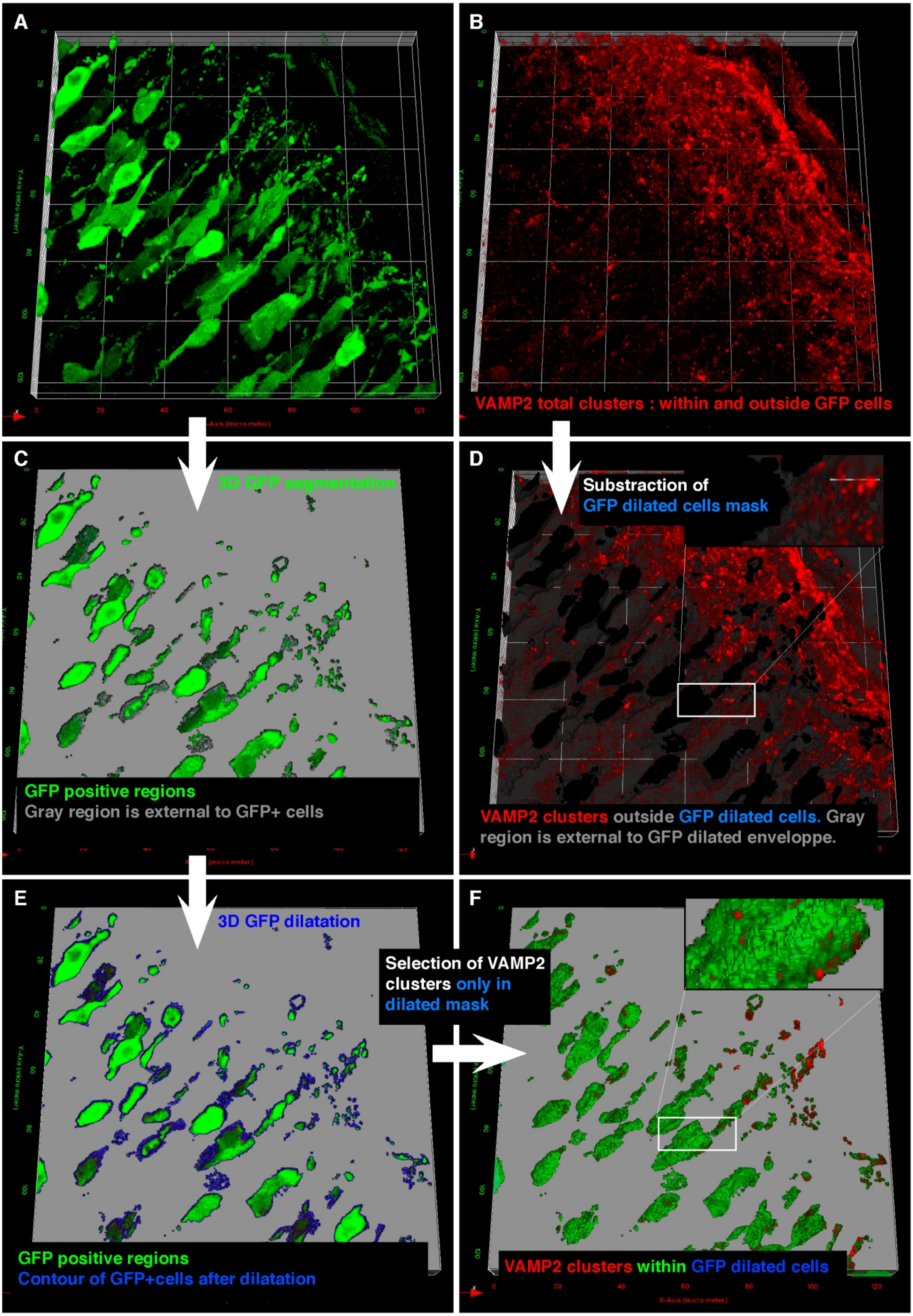
A
B
VAMP2 total clusters : within and outside GFP cells
C
3D GFP segmentation
GFP positive regions
Gray region is external to GFP+ cells
D
Substraction of
GFP dilated cells mask
VAMP2 clusters outside GFP dilated cells. Gray region is external to GFP dilated enveloppe.
E
3D GFP dilatation
GFP positive regions
Contour of GFP+cells after dilatation
F
Selection of VAMP2 clusters only in dilated mask
VAMP2 clusters within GFP dilated cells

Figure S6

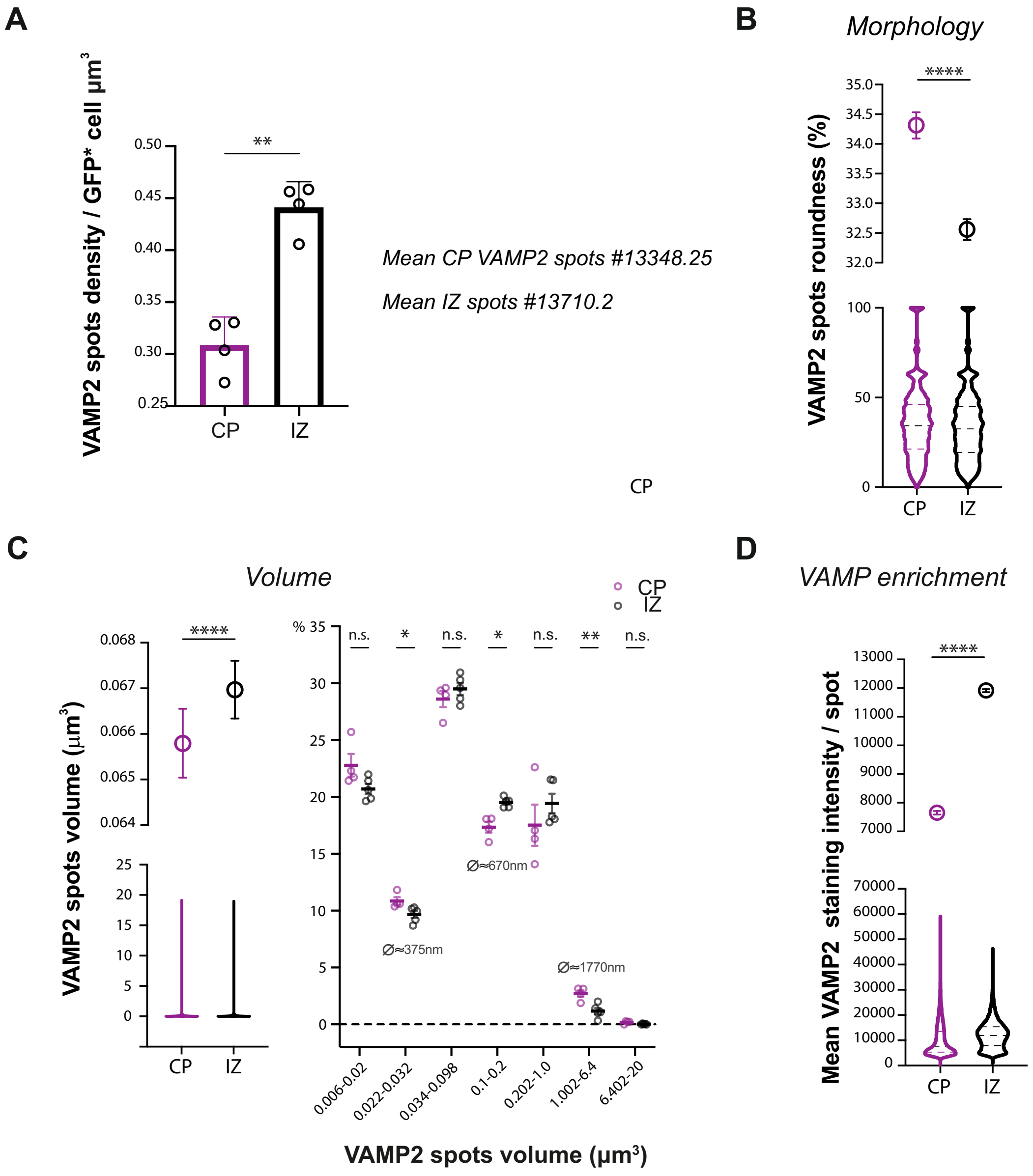
A
VAMP2 spots density / GFP* cell µm³
0.50
0.45
0.40
0.35
0.30
0.25
**
CP
IZ
Mean CP VAMP2 spots #13348.25
Mean IZ spots #13710.2
B
Morphology
VAMP2 spots roundness (%)
35.0
34.5
34.0
33.5
33.0
32.5
32.0
****
100
50
0
CP
IZ
CP
C
Volume
VAMP2 spots volume (µm³)
0.068
0.067
0.066
0.065
0.064
****
25
20
15
10
5
0
CP
IZ
CP
IZ
% 35
30
25
20
15
10
5
0
n.s.
*
n.s.
*
n.s.
**
n.s.
Ø≈375nm
Ø≈670nm
Ø≈1770nm
0.006-0.02
0.022-0.032
0.034-0.098
0.1-0.2
0.202-1.0
1.002-6.4
6.402-20
VAMP2 spots volume (µm³)
D
VAMP enrichment
Mean VAMP2 staining intensity / spot
13000
12000
11000
10000
9000
8000
7000
****
70000
60000
50000
40000
30000
20000
10000
0
CP
IZ

### Figure S7

**Table 1. A summary of migration and synaptogenic phenotypes described for Slitrk1, its mutant forms and RPTPs OE in E13.5-17.5 electroporations**

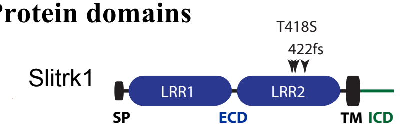


| pCAG constructs | Protein domains | | | | RPTP MeB | Phenotypes | | | | | | | | | | |
|---|---|---|---|---|---|---|---|---|---|---|---|---|---|---|---|---|
| | | | | | | CP1 delay | CP3 delay | MZ/CP presynaptic VAMP2 | | | | IZ stuck cells | IZ presynaptic VAMP2 | | | |
| | LRR1 | LRR2 | TM | ICD | | | | Density | Large spots | Morpho | Intensity | | Density | Large spots | Morpho | Intensity |
| Slitrk1 | + | + | + | + | | + | - | ↑↑ | - | = | ↓ | + | = | + | ⬭ | ↓↓ |
| Slitrk1 ΔECD | - | - | + | + | | - | - | ↑ | - | ⬭ | ↑↑ | - | = | - | ⬭ | = |
| Slitrk1 ΔICD | + | + | + | - | | - | - | ↑↑ | - | ⬭⬭ | ↑↑↑ | - | = | - | ⬭ | = |
| Slitrk1 T418S | + | - | + | + | | +/- * | - | ↑ | - | = | ↑ | + | ↓ * | + | ⬭ | ↓↓ |
| Slitrk1 422fs | + | - | - | - | | - | - | = | - | ○ | ↓ | - | = | - | ○○ | ↓↓ |
| Ptprs | | | | | - | + | - | | | | | - | | | | |
| Slitrk1+Ptprs | + | + | + | + | - | - | - | | | | | + | | | | |
| Ptprd | | | | | + | + | + | | | | | - | | | | |
| Slitrk1+Ptprd | + | + | + | + | + | + | - | | | | | - | | | | |

Absence (-), presence (+), equal to control (=), equal to control and Slitrk1 (+/- * ); ↑ increase and ↑↑ strong increase as compared to control (GFP alone);

↓ decrease and ↓↓ strong decrease as compared to control; ↓ * as compared to Slitrk1; ○ more round and ○○ stronger roundness compared to control.

⬭ elongated and ⬭⬭ increased elongation as compared to control.

**Table 2. Slitrk2 protein domains and disease-associated variants in radial migration**

| pCAG constructs | Protein domains | | | | Phenotype (migration delay) | | |
|---|---|---|---|---|---|---|---|
| | LRR1 | LRR2 | TM | ICD | CP1 | CP2 | CP5 |
| Slitrk2 (WT) | + | + | + | + | - | + | - |
| Slitrk2-V89M | - | + | + | + | - | + | - |
| Slitrk2-L627F | + | + | - | + | + | - | + |

Absence (-), presence (+)
